# Deep learning reveals cancer metastasis and therapeutic antibody targeting in whole body

**DOI:** 10.1101/541862

**Authors:** Chenchen Pan, Oliver Schoppe, Arnaldo Parra-Damas, Ruiyao Cai, Mihail Ivilinov Todorov, Gabor Gondi, Bettina von Neubeck, Alireza Ghasemi, Madita Alice Reimer, Javier Coronel, Boyan K. Garvalov, Bjoern Menze, Reinhard Zeidler, Ali Ertürk

**Affiliations:** Institute for Stroke and Dementia Research, Klinikum der Universität München, Ludwig Maximilian University of Munich (LMU), Munich, Germany; Center for Translational Cancer Research (TranslaTUM) & Department of Computer Science; Graduate School of Neuroscience (GSN), Munich, Germany; Helmholtz Zentrum München, Research Unit Gene Vectors, Munich, Germany; Department of Microvascular Biology and Pathobiology, European Center for Angioscience (ECAS), Medical Faculty Mannheim, University of Heidelberg, Germany; Department for Otorhinolaryngology, Klinikum der Universität München, Munich, Germany; Munich Cluster for Systems Neurology (SyNergy), Munich, Germany; Munich School of Bioengineering, Technical University of Munich, Munich, Germany

## Abstract

Reliable detection of disseminated tumor cells and of the biodistribution of tumor-targeting therapeutic antibodies within the entire body has long been needed to better understand and treat cancer metastasis. Here, we developed an integrated pipeline for automated quantification of cancer metastases and therapeutic antibody targeting, named DeepMACT. First, we enhanced the fluorescent signal of tumor cells more than 100-fold by applying the vDISCO method to image single cancer cells in intact transparent mice. Second, we developed deep learning algorithms for automated quantification of metastases with an accuracy matching human expert manual annotation. Deep learning-based quantifications in a model of spontaneous metastasis using human breast cancer cells allowed us to systematically analyze clinically relevant features such as size, shape, spatial distribution, and the degree to which metastases are targeted by a therapeutic monoclonal antibody in whole mice. DeepMACT can thus considerably improve the discovery of effective therapeutic strategies for metastatic cancer.

**Graphical Abstract:** 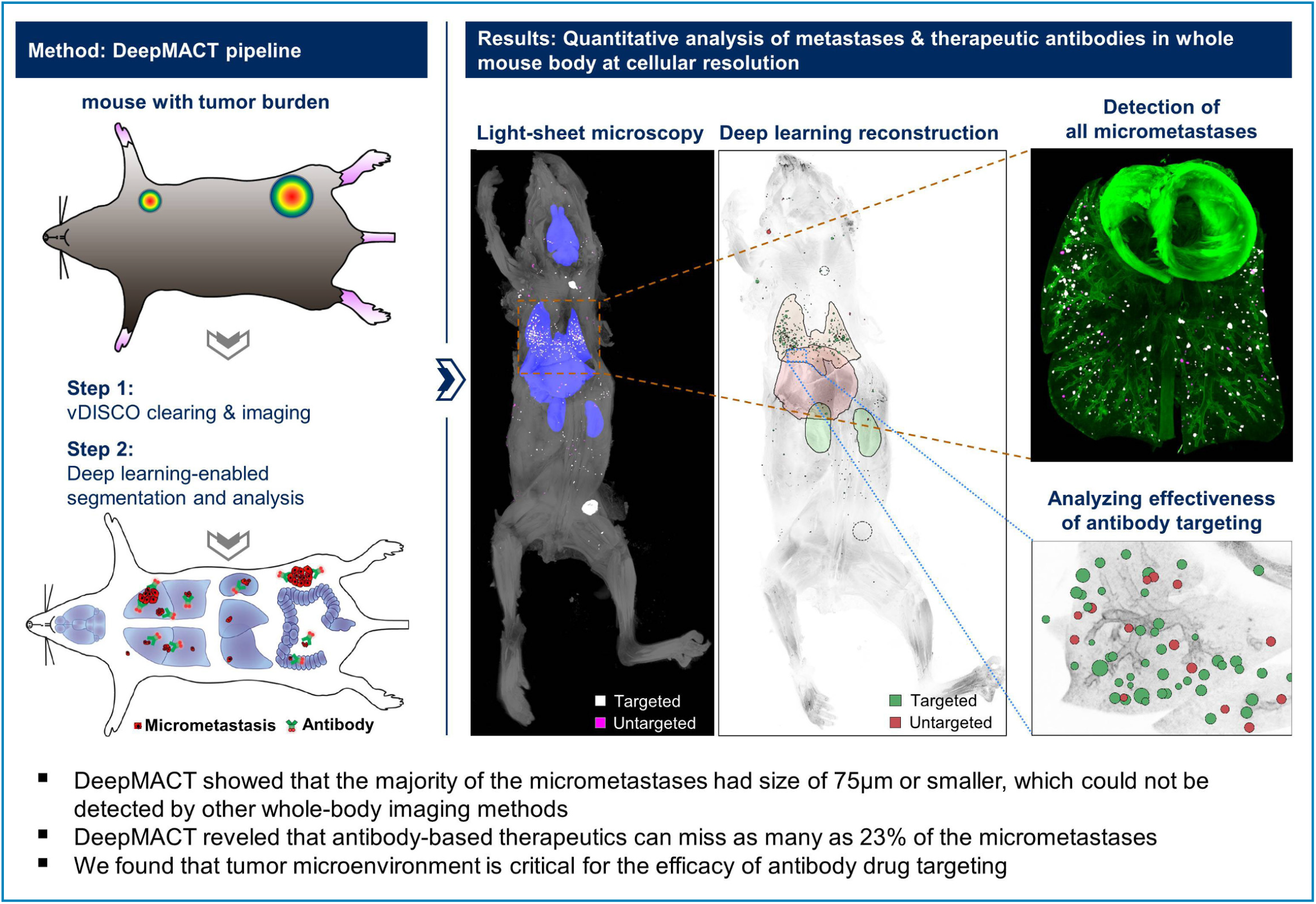

Supplementary Movies and deep learning algorithms of DeepMACT are available at http://discotechnologies.org/DeepMACT/

## INTRODUCTION

Metastasis is a complex process affecting diverse organs^1-3^. As most cancer patients die of metastases at distant sites developing from disseminated tumor cells with primary or acquired resistance to therapy, a comprehensive and unbiased detection of disseminated tumor cells and tumor targeting drugs within the entire body at the single cell level is crucial^4^. Such technology would help to explore mechanisms affecting tumor metastasis and drug targeting much more reliably, hence substantially contributing to the development of improved therapeutics. So far, such efforts have been hampered by the lack of 1) imaging technologies to reliably detect single cancer cells in intact mouse bodies, and 2) algorithms to quickly and accurately quantify large-scale imaging data. Here, we developed an analysis pipeline that allows us to efficiently solve these limitations.

First, we built upon recently developed whole mouse clearing methods^5-8^ to address the imaging problem. Typically, fluorescent labeling of cancer cells *in vitro* or *in vivo* is achieved by endogenous expression of fluorescent proteins such as GFP, YFP, and mCherry, which emit light in the visible spectrum. However, many tissues in the body show high autofluorescence in this range^9,10^, which hinders reliable detection of single cancer cells or small cell clusters in intact mouse bodies based on their endogenous fluorescent signal. To circumvent this problem, we chose to implement the vDISCO technology^8^, which amplifies the signal of fluorescent proteins of cancer cells more than 100-fold, enabling reliable imaging not only of large metastases but also micrometastases down to single cells throughout the entire body.

Second, systematic analysis of metastasis in whole adult mouse bodies requires quantitative information such as location, size and shape of all individual metastases. Manual detection and segmentation of numerous metastases in highly resolved full body scans is an extremely laborious task that may take several months per mouse for an expert annotator. In addition, automation by filter-based 3D object detectors is not reliable as different body tissues have different levels of contrast^5^, causing a high rate of false positive and false negative cancer cell detection. Recent studies have started to show the high-efficacy of deep learning-based analysis of biomedical images, as well as pre-clinical studies, compared to filter-based or manual segmentation methods^11-17^. To enable automated, robust, and fast mapping of all tumor cells in transparent mice, we developed an efficient deep learning approach based on convolutional neural networks (CNNs) and optimized it for the vDISCO imaging data and tumor cell distribution patterns.

Together, resolving these two bottlenecks allowed us to build an integrated, highly automated pipeline for analysis of metastasis and tumor-targeting therapeutics, which we named DeepMACT (Deep learning-enabled Metastasis Analysis in Cleared Tissue). Using DeepMACT, we detected cancer metastasis and therapeutic antibody targeting at the single cell level in entire mouse bodies, including many metastases previously overlooked by human annotators. As a scalable, easily accessible, fast, and cost-efficient method, DeepMACT enables a wide range of studies on cancer metastasis and therapeutic strategies. To facilitate adoption, the protocols for clearing and imaging, as well as the deep learning algorithm, the training data, and the trained model are freely available online for adaptations to address diverse questions in pre-clinical and clinical research.

## RESULTS

Focusing on a clinically relevant tumor model, we transplanted human MDA-MB-231 mammary carcinoma cells, expressing mCherry and firefly luciferase, into the mammary fat pad of NOD *scid* gamma (NSG) mice and allowed the tumors to grow and metastasize for 6-10 weeks (**Figure 1A**)^18-20^. Furthermore, we injected the fluorescently-tagged 6A10 therapeutic antibody that has been shown to reduce tumor burden in this model^19,20^. To investigate cancer metastasis and therapeutic antibody targeting in whole mouse bodies at the single cell level, we developed DeepMACT. In short, we transcardially perfused the animals using standard PFA fixation and applied the vDISCO method to boost the fluorescent signal of tumor cells in transparent mice. After light-sheet microscopy, the 3D image stacks of whole mouse bodies were analyzed using deep learning algorithms. The DeepMACT pipeline consists of 1) vDISCO panoptic imaging of transparent mice at the single cell level and 2) deep learning-based analysis of cancer metastasis and antibody drug targeting (**Figure 1B**).

**Figure 1.**
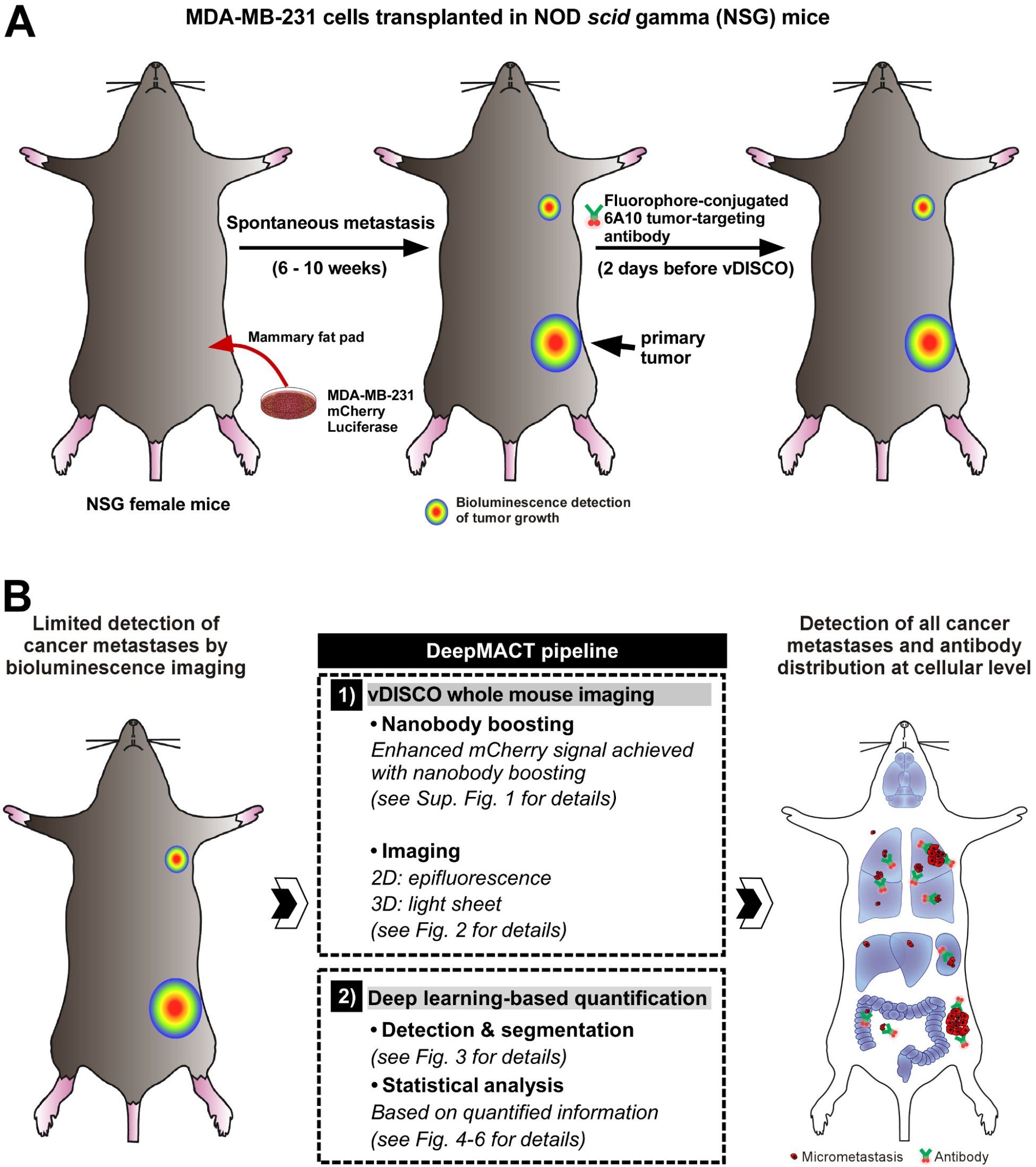
Experimental design and schematic of DeepMACT pipeline for analysis of cancer metastases and antibody drug targeting at single cell level. (**A**) Illustration of the experimental workflow for tumor transplantation and antibody application. (**B**) Steps of the Deep-MACT pipeline on whole mouse bodies. First, the whole mice are fixed and processed with the vDISCO protocol to amplify the fluorescent signal of cancer cells. Whole transparent mice are subsequently imaged using light-sheet mi-croscopy from head to toe at single cell resolution. Light-sheet images are assembled into a complete 3D image of the mouse. Next, convolutional neural networks are trained to identify and segment all micrometastases in the fluorescence signal. The trained algorithms are then applied to 3D images to detect cancer metastases and an antibody-based drug targeting in whole transparent mice at single cell level.

### DeepMACT step 1: vDISCO imaging of cancer metastases at cellular resolution in intact mice

We previously developed the vDISCO technology to image single cells in mouse bodies through intact bones and skin^8^. The vDISCO method utilizes bright fluorescent dyes tagged to nanobodies to boost the signal of fluorescent proteins expressed in the target cells. Here, we first applied vDISCO to increase the fluorescence signal of mCherry-expressing cancer cells. Boosting the tumor cell fluorescence with anti-mCherry nanobodies conjugated to Atto-594 or Atto-647N dyes, we found that nanoboosters can enhance the signal quality of cancer cells over 100 times compared to imaging the endogenous mCherry signal (**Figure S1**). Owing to this significant enhancement in signal contrast, we could readily detect single cancer cells buried in centimeters-thick mouse bodies, e.g., in deep brain regions through the intact skull (**Figure S1F**, yellow arrowhead). To confirm the specificity of vDISCO boosting of mCherry expressing cancer cells, we performed the following experiments: 1) we stained control mice without a tumor transplant, thereby lacking mCherry expression, and found no labeling in any of the analyzed organs; 2) we analyzed the primary tumors and lung metastases from the vDISCO-processed mouse bodies by staining them using a specific anti-luciferase antibody, which confirmed that endogenous mCherry fluorescence co-localized with both the nanobooster and the luciferase signals; and 3) we confirmed the colocalization of nanoboosters with mCherry expressing single tumor cells (**Figure S2**).

Since the detection of smaller-sized tumor cell clusters, which may represent dormant cancer cells or incipient metastatic nodules, is critical, we next tested if vDISCO allows imaging cancer metastases in whole-mouse bodies at the single cell level. In order to compare our approach to conventional methods, we also acquired bioluminescence images of mice before applying DeepMACT. In line with previous findings^18^, we detected the earliest large metastasis of transplanted MDA-MB-231 cells at the axillary lymph node of mice by bioluminescence (**Figure 2A, Figure S3**). However, bioluminescence imaging did not reveal any detailed information such as size or shape and failed to show the presence of micrometastases.

**Figure 2.**
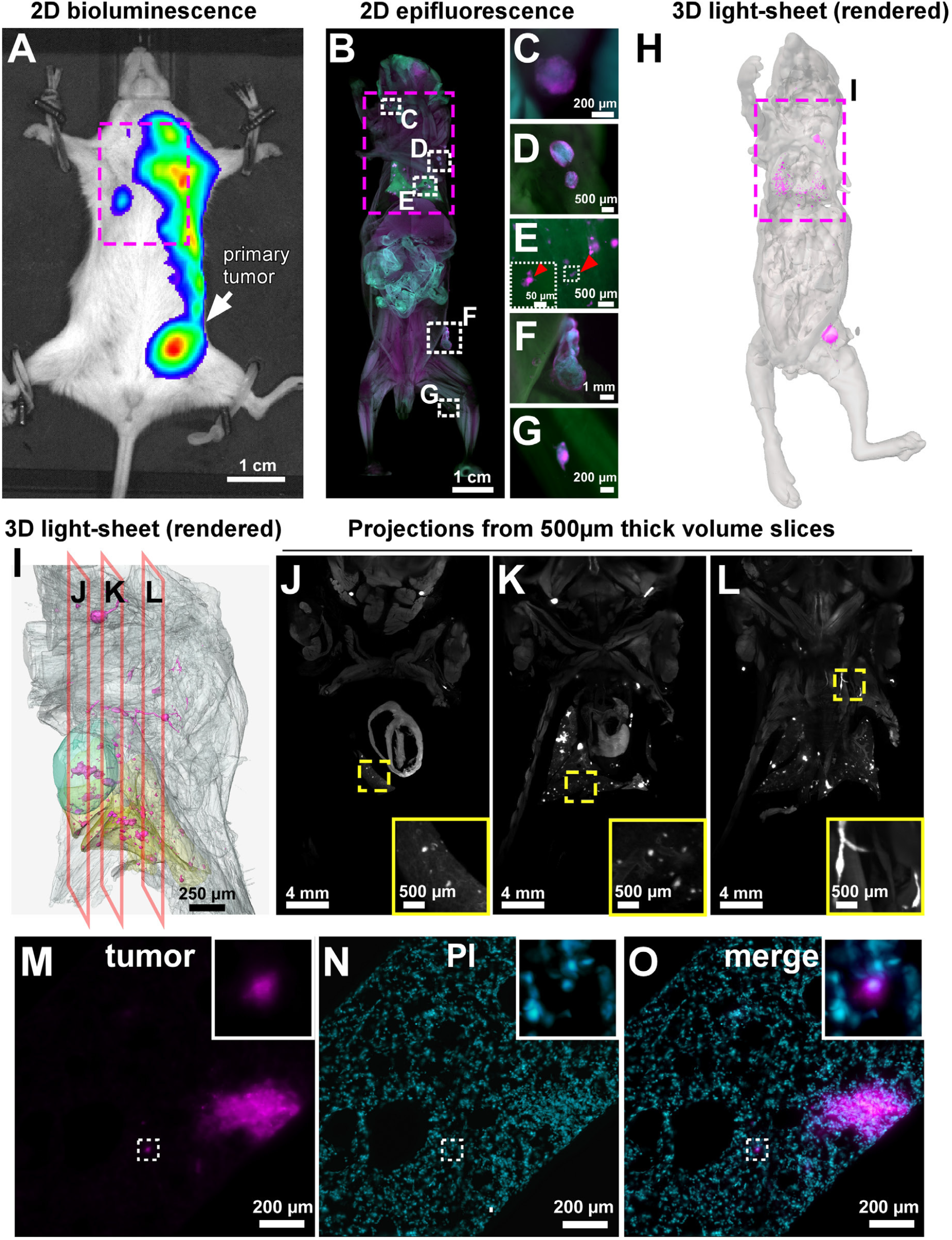
DeepMACT step 1: vDISCO visualization of metastases in an intact whole mouse body. A) Bioluminescence image of a tumor-bearing mouse before vDISCO. (**B**-**G**) Epifluorescence images of the same mouse after vDISCO show metastases (magenta) in greater detail compared to bioluminescence, including small micrometastases that can be readily detected in the lungs (**E**, red arrowhead) and in the leg (**G**), in addition to the primary tumor (**F**) and major metastases (**C** and **D**) that are also visible in bioluminescence as bulk signal (A). (**H**) 3D visualization of the intact transparent body of the same mouse, imaged by light-sheet microscopy. (**I**) Lateral views of the 3D segmentation obtained from the light-sheet imaging data corresponding to the magenta-boxed region indicated in (A, B, and H). For simplicity, only a few organs are segmented: the heart (cyan) and the lungs (yellow); the mouse body is shown in transparent gray and the metastases are in magenta. (**J**-**L**) Original light-sheet microscopy data (500 µm projections) showing tumors from the sagittal planes indicated in (J). (**M**-**O**) High resolution light-sheet microscopy images (single planes) showing single tumor cells (magenta) and nuclei (labeled with propidium iodide, PI; cyan) revealed by vDISCO. See Figure S1-S4, Movie S1 and S2.

After bioluminescence assessment, we applied vDISCO using anti-mCherry nanoboosters conjugated to Atto-647N and imaged the whole-mouse bodies first using epifluorescence in 2D (**Figure 2B-G**), then using light-sheet microscopy in 3D (**Figure 2H-O**). In epifluorescence, we could readily see both the primary tumor (**Figure 2F**) and the major metastases at the axillary lymph node (**Figure 2D**), which were also detected by bioluminescence imaging (**Figure 2A**), albeit as a bulk signal, lacking information on real size and shape. By contrast, our approach allowed the visualization of several micrometastases in the lungs with conventional epifluorescence imaging, which were not visible in bioluminescence (compare the magenta marked regions in **Figure 2A** with **Figure 2B,** and red arrowheads in **Figure 2E**; more examples shown in **Figure S3**). Thus, vDISCO followed by epifluorescence imaging, which can be completed within minutes, already provided greater details and sensitivity compared to bioluminescence imaging. Next, we imaged entire transparent mice using a light-sheet microscope^8^ at cellular resolution in 3D to detect micrometastases throughout the body (**Figure 2H**). In the chest area, we could see various metastases not only in the lungs (yellow segmented region in **Figure 2I**) and lymph nodes, but also at the base of the neck and surrounding tissues (**Figure 2J-L** and **Movie S1**). Importantly, light-sheet microscopy scanning allowed us to image micrometastases down to the single cell level in the intact mouse body. An example of a single-cell metastasis is shown in the dashed box in **Figure 2M** (also see **Movie S2**) as verified by nuclear staining with PI (**Figure 2N-O)**. Thus, our approach allows, for the first time, to image micrometastases in intact mice in 3D down to single cell resolution.

### DeepMACT step 2: Deep learning for detection and quantification of metastases

We developed a novel deep learning-based approach to detect and segment all cancer cells in whole mouse bodies. This framework solves the 3D task of detecting and segmenting metastases in volumetric scans with CNNs that process 2D projections of small sub-volumes (**Figure 3A**). In brief, we first derived three 2D maximum intensity projections (aligned with the x-, y-, and z-axes) for each sub-volume in order to increase the signal-to-noise ratios (SNR). We fed the resulting projections to the CNN and obtained 2D probability maps, in which each pixel value represents the estimated probability that this pixel identifies a metastasis under the given projection. We then reconstructed a 3D segmentation from the three projections observing increased reliability in detecting true positive metastases while safely ignoring non-metastatic tissue that would be false positives in the individual projections. For example, in **Figure 3B**, the green arrows show successful detection of a real metastasis and the red arrows show successful ignoring of a structure that could be mistaken for a metastasis from a single 2D projection. This approach was highly effective in detecting and segmenting metastases in the imaging data, yielding a binary mask for all metastases in the body.

**Figure 3.**
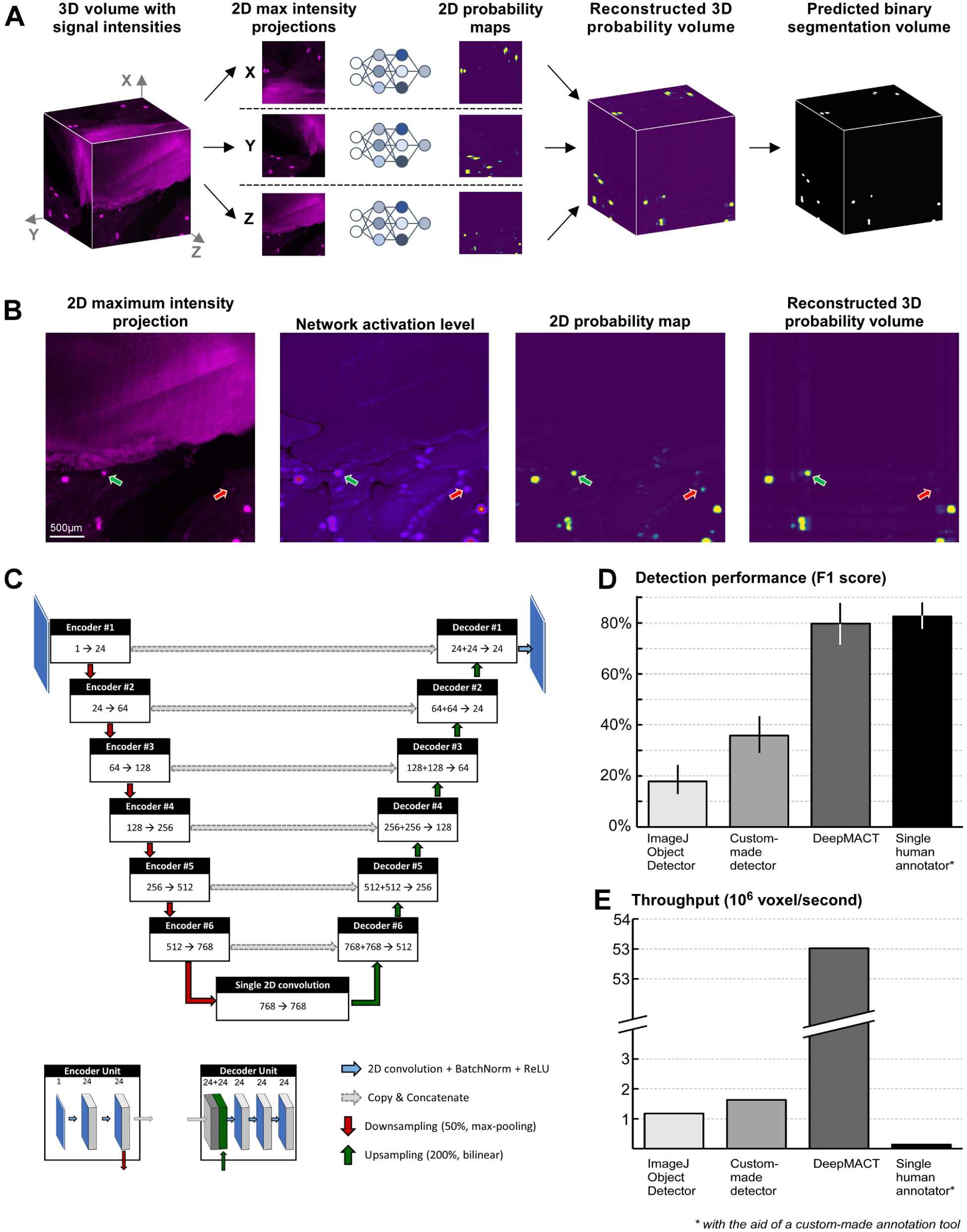
DeepMACT step 2: Architecture and performance of the deep learning algorithm. (**A**) Representation of the deep learning inference workflow to efficiently derive 3D detection and segmentation exploiting three 2D computational operations. (**B**) Visualization of the computational stages; the green arrow shows successful detection of a metastasis, the red arrow shows elimination of a potential false positive detection after 3D recombination. (**C**) High-level representation of the network architecture with an encoding and a decoding path. (**D**-**E**) Comparison of our deep learning pipeline to alternative automated methods and manual segmentation by a human expert in terms of performance (**D**) and speed (**E**).

The core of the architecture makes use of CNNs (**Figure 3C**), structurally similar to the established U-net^21^, which learn to distinguish metastases from the background signal. This is achieved by using a deep stack of encoding units, which detect characteristic cancer features, and a corresponding stack of decoding units, which segment each metastasis at pixel-level. Each encoding unit performs two convolutions, extracting information about the environment for each pixel and representing that information in a third dimension - the feature channels. Before being passed on to the next encoding unit, the image is spatially down-sampled. Together, this means that the neural network is steadily increasing the feature channels and steadily decreasing the spatial resolution, enforcing the network to learn even more abstract representations of the data (i.e., features) in the deeper layers, before mapping the information relevant to cancer cells back to the original resolution in the decoding upward path. This happens by up-sampling the abstract, low-resolution information from lower layers and concatenating it with the less abstract, but higher-resolution information from the encoding path via skip connections (some exemplary visualizations of the computational stages are presented in **Figure S4A-C**).

To assess the reliability of our automated deep learning architecture, we applied it to a fresh test set of a full-body scan, which was neither used for training the CNNs nor to optimize hyperparameters. The data sets were manually annotated by human experts and any disagreements between experts were jointly reviewed and discussed in order to derive a refined, commonly agreed reference annotation (see methods for details). We then systematically compared the performance of our deep learning approach to that of established detection methods as well as the performance of a single human annotator, calculating F1-score (also known as Dice score), a common performance measure based on both the metastasis detection rate (recall) and false positive rate (precision).

We found that DeepMACT reached an F1-score of 80%, outperforming existing filter-based detectors by a large margin and coming very close to the level of a single human annotator with an F1-score of 83% (**Figure 3D**). The slightly higher F1-score of a single human annotator is mainly driven by the high precision. However, the human annotator missed around 29% of all micrometastases (examples are shown in **Figure S4D-F**) and detecting those false negatives would require a repetitive and very laborious re-analysis of the whole animal scans – several months of human work time. On the other hand, the F1-score of DeepMACT is a result of a balance between precision and recall, which can be freely adjusted via the model’s threshold. For DeepMACT, we can increase detection rate (recall) over 95%. While this also increases the false-positive rate, correcting the false positive data requires only a review of marked signals by a human annotator, which we completed within hours for the data of this study (an example is shown in **Figure S4G-I**). A more detailed analysis on the trade-off between precision and recall is shown in **Figure S4J**. Notably, DeepMACT could detect micrometastases about 30 times faster than filter-based detectors and over 300 times faster than a human annotator (**Figure 3E**) who was already supported by a dedicated and interactive software, custom-built for this task and these data; without annotation software, the human manual annotation would be estimated to take several months for a single mouse. Thus, DeepMACT can complete months to years of human labor in within hours without compromising on segmentation quality.

### DeepMACT detects micrometastases at the cellular level

After establishing the DeepMACT pipeline, we used it to analyze whole mouse bodies. In addition to the primary tumor and the macrometastasis in the axillary lymph node, we could detect hundreds of micrometastases of varying sizes throughout the body, especially in the lungs (**Figure 4A-B)**. Overall, DeepMACT identified 520 micrometastases throughout the entire body in this particular mouse, of which there were 306 in the lungs, 5 in the liver, 1 in the left kidney and 208 in the rest of the body (**Figure 4C**). We found that micrometastases are mostly located in the inner tissue layers (about 1 cm depth from the surface), as shown by color-coding in **Figure 4D,** making them extra difficult to detect by other methods. To analyze the spatial distribution with regard to the lung anatomy, we registered all 306 lung micrometastases to the mouse lung lobes. We found that micrometastases were evenly distributed in all lobes (**Figure 4E,F**). Interestingly, the micrometastases were randomly distributed throughout the lungs regardless of their size, suggesting independent colonization at multiple sites. Furthermore, we quantified the size and relative location of all micrometastases in the entire body (**Figure 4G-L**). While 79% of micrometastases were within 1 mm to the nearest neighboring micrometastasis, we also found highly isolated micrometastases as distant as 9.3 mm apart from their nearest neighbor (**Figure 4G**). Importantly, we found a large number of micrometastases with only a few hundred cells (**Figure 4H**) and diameters less than 50-100 µm (**Figure 4I**), which would be very difficult to detect in intact mice by other methods. Comparing the micrometastases in the lungs with the torso, we found that the tumor burden in the lungs was more than a hundred times higher (**Figure 4J**). Also, micrometastases in the lungs were, on average, 30% larger in diameter (**Figure 4K**) containing more than twice as many cells per metastasis as micrometastases in the rest of the torso (**Figure 4L**). In sum, our pipeline is the first to enable quantitative analyses of whole-body scans at cellular resolution, yielding important insights into understanding the metastatic process.

**Figure 4.**
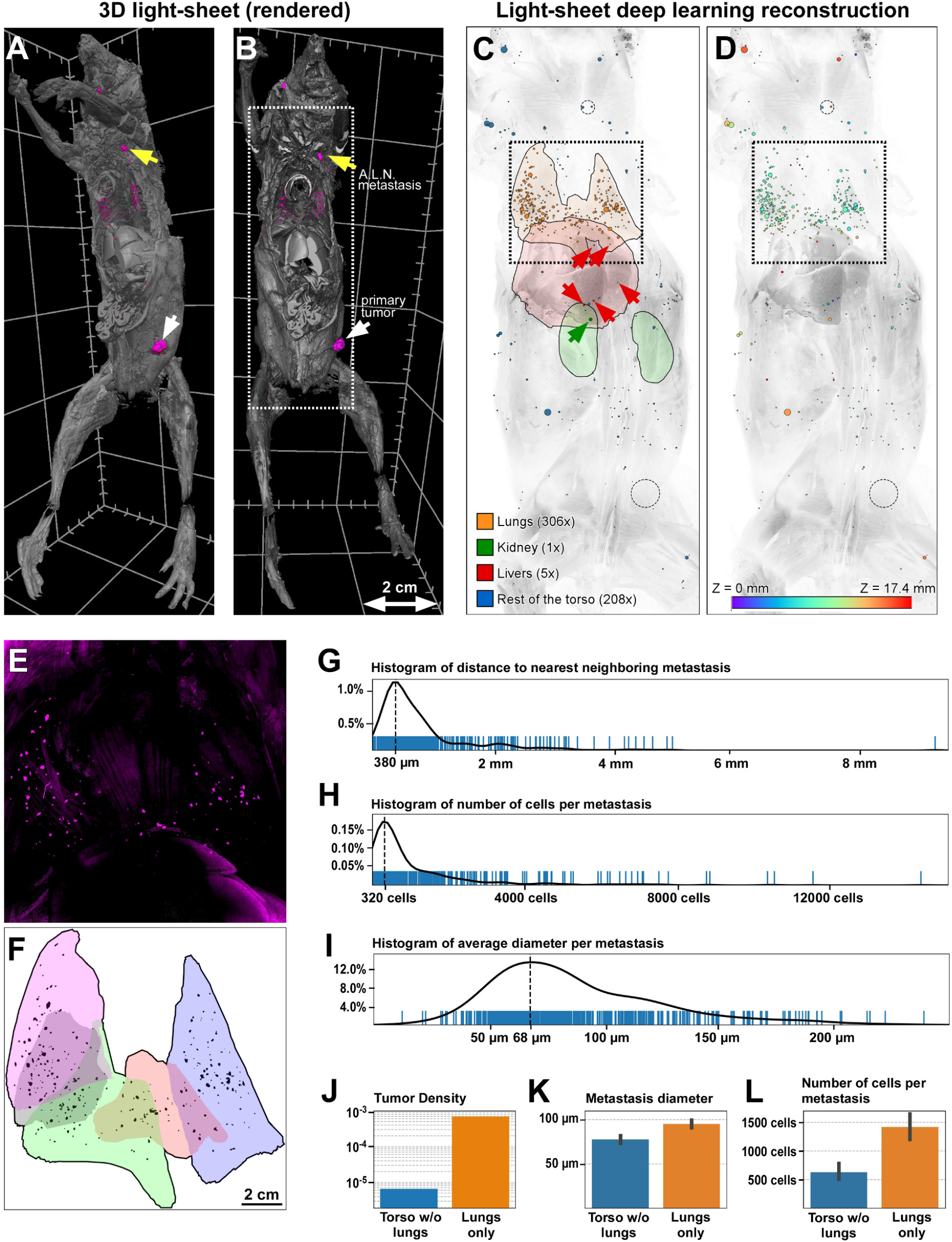
Deep learning-based detection and segmentation enables quantitative analysis at the level of individual metastases. (**A**,**B**) 3D rendering of the entire mouse after light-sheet microscopy imaging in ventral and lateral views, respectively. Metastases in the mouse body are shown in magenta. The white arrow indicates the primary tumor and the yellow arrow indicates metastases in the axillary lymph node (A.L.N.). (**C**,**D**) Deep learning reconstructions of all detected metastases (A.L.N. and primary tumor indicated with dashed circles) color-coded by organ (**C**) and depth along z-axis (**D**), cropped to white box of (A) to show higher level of detail. (**E**,**F**) Detailed view of metastases in the lung region (corresponding to the black box in C) in a maximum intensity projection of a 3D light-sheet scan (**E**) and projection of 3D deep learning-based detection, with metastases registered to individual lung lobes (shown in different colors) (**F**). (**G**-**L**) Deep learning-based distributions; blue bars show individual metastases, the black line shows the Gaussian kernel density estimation. (**G**) 3D distance to nearest neighboring metastasis. (**H**) Cell count estimation per metastasis. (**I**) Metastasis diameter averaged in 3D space. (**J**-**L**) Quantitative comparison between metastases in the lungs and the rest of the torso; bars indicate 95%-confidence intervals. (**J**) Tumor density as share of metastatic tissue of the entire volume. (**K**) Metastasis diameter averaged in 3D space. (**L**) Cell count estimation per metastasis.

### DeepMACT reveals therapeutic antibody targeting at the cellular level

A number of tumor-targeting monoclonal antibodies have become part of the standard treatment for various solid and hematological malignancies and many more are in early or late stages of clinical development^22,23^. However, so far there has been no methodology to determine the distribution of therapeutic antibodies in the entire body at cellular resolution. Here, we used DeepMACT to assess biodistribution of the therapeutic monoclonal antibody 6A10 directed against human carbonic anhydrase XII (CA12)^19,20,24^. CA12 is overexpressed in various types of cancers and blocking its activity with the antibody 6A10 reduces tumor growth^19^ and increases the sensitivity of tumor to chemotherapy^20^. We intravenously injected 20 µg of 6A10 conjugated to Alexa-568 (with tumor signal boosted with Atto-647N) nine weeks after transplantation of MDA-MB-231 cells and perfused the mice two days after the antibody injection for whole-body analysis. Because Alexa-568 excitation/emission spectra overlap with the endogenous mCherry signal of the transplanted cancer cells, we confirmed that the vDISCO pipeline eliminates all the signal from endogenously expressed mCherry^8^ (**Figure S5**).

We first acquired 2D images with epifluorescence microscopy and observed an accumulation of the 6A10 antibody at the primary tumor (**Figure 5A,E**; tumor shown in magenta, therapeutic antibody in cyan) and the metastases at the axillary lymph node (**Figure 5A,B**). Focusing on the lungs, we detected micrometastases that were targeted by the 6A10 antibody (**Figure 5C**, white arrow) and others that were not (**Figure 5D**, yellow arrow). Acquiring 3D scans with light-sheet microscopy, we assessed the complete biodistribution of the therapeutic antibody and micrometastases in the whole body at cellular resolution (**Figure 5F-H, Movie S3)**. The axillary lymph node metastases and the micrometastases in the lungs are shown in **Figure 5F**. Analyzing the signal of individual micrometastases and the 6A10 antibody by light-sheet microscopy in 3D, we could evaluate the efficiency of antibody drug targeting for even the smallest micrometastases (**Figure 5G**, white arrowhead). We also verified the targeting of micrometastases by the 6A10 antibody in various organs such as lungs and kidney, using confocal microscopy (**Figure S6**).

**Figure 5.**
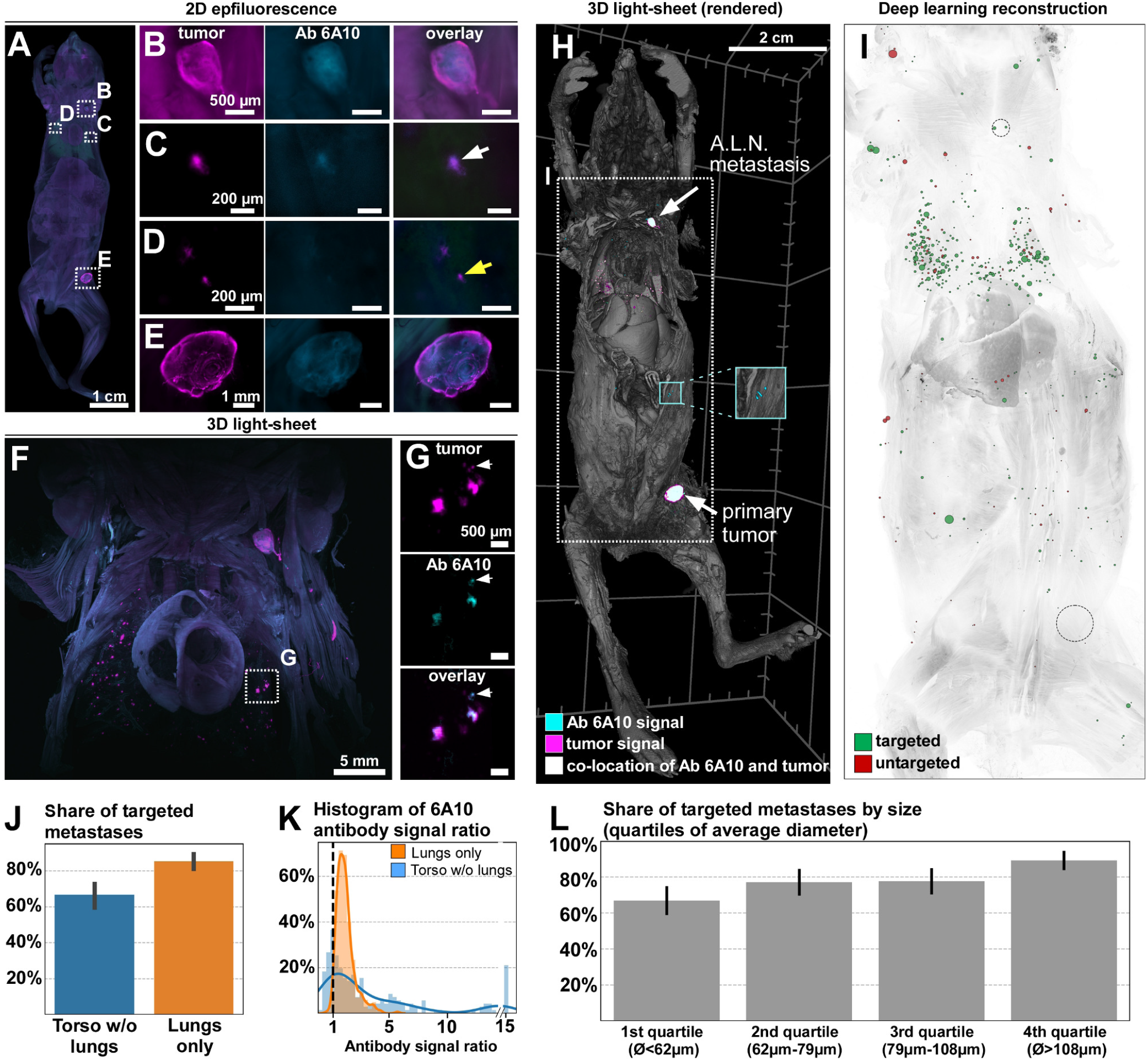
The DeepMACT pipeline enables quantitative analysis of drug delivery effectiveness at the level of single metastases. (**A**) Epifluorescence images of a processed mouse show details (**B**-**E**) of both tumor metastases (boosted with Alexa647 nanobody, shown in magenta) and 6A10 antibody (conjugated with Alexa568, shown in cyan) distributions and their overlay. While most of the micrometastases are targeted by the antibody (**C**, white arrow), there are some that are not (**D**, yellow arrow). (**F**) Full body 3D light-sheet scan, cropped to the chest region, shows the distributions of metastases (magenta) and antibody (cyan). (**G**) Detailed view of the boxed region in (F) showing very small micrometastases targeted by the therapeutic antibody (white arrows). (**H**) 3D rendering of a whole mouse body light-sheet scan showing the tumor signal in magenta, the 6A10 antibody signal in cyan (co-localization of the signals is shown in white). The cyan inlet shows an example of off-target accumulation of the 6A10 antibody. (**I**) Deep learning-based reconstruction of the animal in (H) showing targeted metastases in green and untargeted metasases in red; the dashed circles represent the primary tumor A.L.N metastases. (**J**) Comparison of the share of targeted metastases in the lungs versus the rest of torso. (**K**) Comparison of the distributions of 6A10 antibody signal ratio (signal strength in metastasis versus local surrounding; see the methods for further details) per metastasis in the lungs versus the rest of torso. (**L**) Share of targeted metastases as a function of their size (split into quartiles of average metastasis diameter).

Next, we used DeepMACT to systematically assess and quantify the effectiveness of antibody drug targeting in whole animals at the cellular level (**Figure 5I**). While overall 77% of all metastases were targeted by the antibody, we found that significantly more micrometastases (*p* < 0.001, two-sided *t*-test) were targeted in the lungs (85%) as compared to the rest of the body (66%) (**Figure 5J, Movies S3 and S4**). To further assess the efficacy of drug targeting for micrometastases in the lung versus the rest of the body, we assessed the antibody concentration by quantifying the antibody signal contrast (relative signal strength versus local surrounding; see methods for details) (**Figure 5K**). Metastases in the lungs generally tend to have a higher antibody signal ratio. This is in line with the higher share of targeted metastases. Also, the antibody signal ratio is much more narrowly distributed compared with micrometastases outside the lungs. The lower average and wider distribution of antibody signal ratio in the micrometastases in the rest of the body indicates that there is a substantially higher variance in the antibody targeting to the cells of those micrometastases. While some are very strongly targeted, many others are not targeted at all. Also, the largest quartile of micrometastases was significantly more likely (*p* < 0.001, two-sided *t*-test) targeted (88%) than the smallest quartile (67%) (**Figure 5L**). We also identified various off-target binding sites throughout the body, i.e. binding of the therapeutic antibody to mouse tissues, which is presumably due to unspecific binding since 6A10 does not bind to murine CA12 (cyan inlet in **Figure 5H**). Overall, these data demonstrate that DeepMACT provides a powerful platform to track the biodistribution of therapeutic antibodies along with micrometastases in intact mouse bodies. Thus, it represents the first methodology that allows quantitative analysis of the efficacy of antibody-based drug targeting in the whole body at cellular resolution.

### Exploring potential mechanisms of antibody drug targeting

The above results demonstrated that antibody-based drugs, which are the basis of many targeted/ personalized treatments, may miss as many as 23% of the micrometastases. Next, we aimed to explore potential mechanisms that might explain this failure. We first hypothesized that the efficacy of targeting of micrometastases might depend on the presence of nearby blood supply transporting the therapeutic antibody. To explore if the vascularization of defined tissue regions can have an effect on antibody drug targeting, we performed lectin labeling of vessels in the lungs, where most of the micrometastases are located. Analyzing diverse micrometastases of different sizes, we found that each of them had blood vessels as close as 1-6 µm (**Figure 6A,B**). This distance is smaller than even a single cell diameter (^∼^ 10 µm) suggesting that the proximity of blood vessels could not be the major reason for the lack of antibody drug targeting^25^.

**Figure 6.**
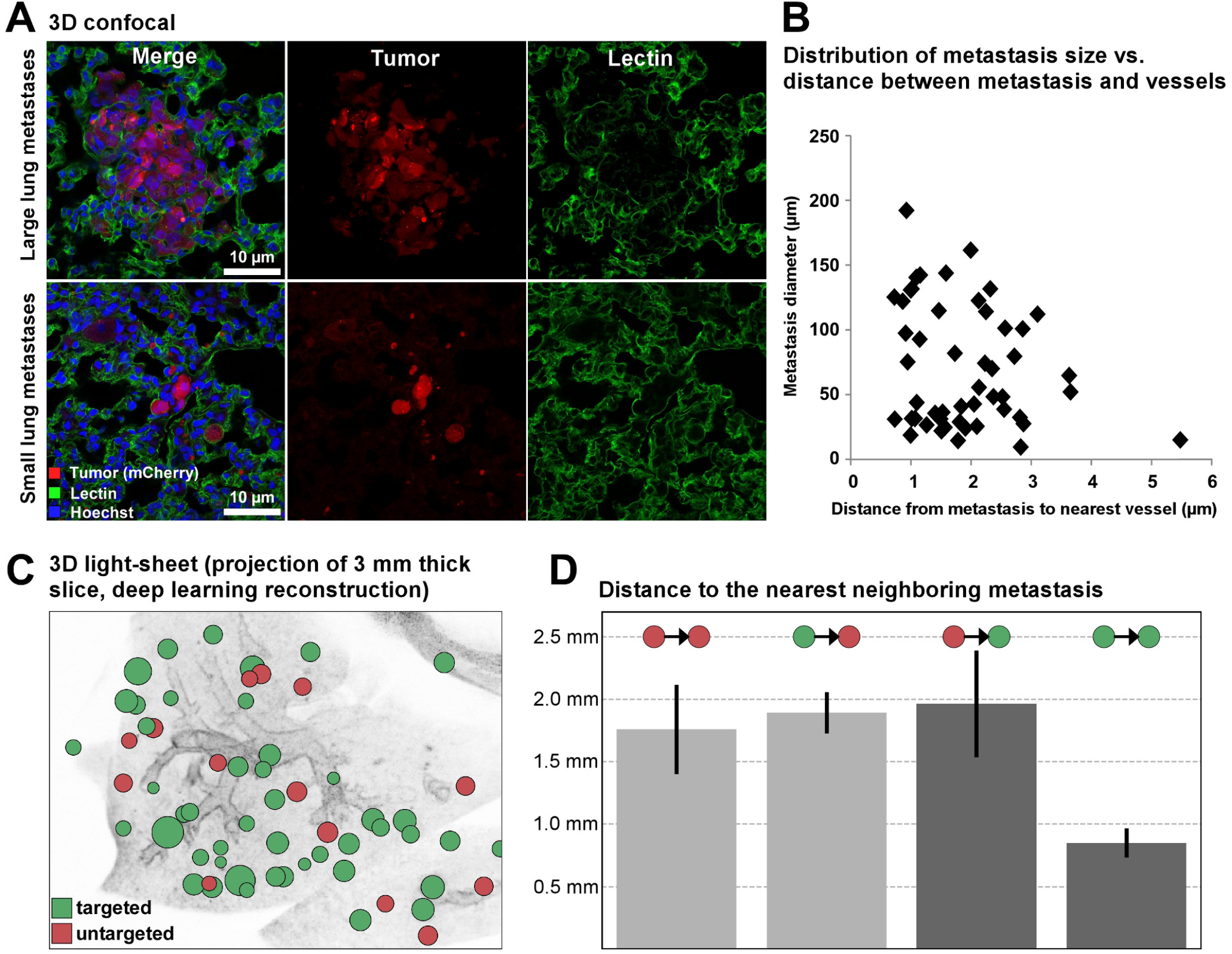
Potential mechanisms of tumor targeting by therapeutic antibody. (**A**) Confocal images of a large and a small metastasis (less than 5 cancer cells) in lungs labeled with Lectin (green) and Hoechst (blue). (**B**) Distribution of metastasis size and distance to the nearest vessel, showing that most of the metastases are close to vessels (distance less than 6 µm) (n=50). (**C**) Deep learning-based reconstruction of lung metastases with and without 6A10 antibody targeting. (**D**) Deep learning-based quantification of distance between metastases and their nearest neighbor, showing local clustering of targeted and untargeted metastases (see the Methods for further details)

Next, we hypothesized that the tumor micro-environment at the sites of metastases could be related to the efficiency of targeting. If so, we would expect a non-random spatial distribution of targeted and untargeted metastasis on a local scale. To address this, we turned to DeepMACT and assessed local clustering of micrometastases targeted by the antibody. We quantified the distances between micrometastases and their nearest neighbor for all micrometastases in the entire body, differentiating between targeted and untargeted nearest neighbors. The distance between two neighboring metastases is significantly smaller for two targeted metastases (about 0.8 mm) than for two untargeted or a mixed pair of an untargeted and a targeted metastasis (consistently at about 1.7-2.0 mm) (**Figure 6C,D**). Importantly, the average distance from an untargeted to the nearest targeted metastasis is significantly larger than from a targeted one. This would not be expected in a random distribution and indicates a clustering on a local scale. Thus, these analyses suggest the existence of factors in tumor microenvironments on a millimeter scale influencing the efficacy of antibody drug targeting.

## DISCUSSION

Unbiased detection of cancer metastasis and the biodistribution of tumor-targeting therapeutics at the single cell level would substantially accelerate pre-clinical cancer research. Towards this goal, we capitalized on a powerful whole-body clearing and imaging method combined with deep learning-based analysis, enabling us to visualize and analyze cancer metastasis in intact transparent mouse bodies. The resulting DeepMACT workflow is a straightforward methodology for systemic analysis of micrometastases and therapeutic antibody drug distribution in whole mouse bodies at cellular resolution within days, a task that would otherwise take several months to years of human labor. Notably, DeepMACT can readily be applied in diverse labs without the need for highly specialized equipment as we provide detailed instructions on VDISCO whole body clearing and imaging, and make the algorithms, the training data, and the trained model publicly available (see methods). Thus, DeepMACT-based evaluation of whole mouse bodies instead of selected tissues/organs at the single cell level can foster the translation of new therapies into the clinic much more efficiently than traditional methods.

### DeepMACT technology

Here we set out make use of recent technologies that are providing scalable and unbiased histological assessment of entire biological specimens. Most whole-body clearing and imaging studies have so far relied on visualization of endogenous fluorescent signal, which did not allow imaging and quantification in intact transparent mice^5,26^. To overcome this, here we adopted the vDISCO whole mouse transparency technology, as it amplifies the signal of cancer cells more than 100 times, ensuring reliable detection of single cells through intact bones and skin^8^. Because vDISCO employs nanobody enhancement of the endogenous fluorescent signal, currently up to 21 types of fluorescent proteins can be boosted with available nanobodies. In addition, conjugation of existing nanobodies with fluorescent dyes at diverse spectra, including those in the near infrared range would help to generate more options for multiplex experiments including imaging of more than one type of fluorescently labeled cell along with conjugated therapeutic antibodies.

Secondly, we developed a highly efficient deep learning architecture based on U-net like CNNs exploiting 2D maximum-intensity projections with high SNR to reliably detect metastases in 3D. Deep learning-based detection not only serves the purpose of automation, but is also provides a very effective tool in finding metastases that would be easily overlooked by humans. In our data, a single human annotator missed around 29% of all metastases. This is in line with previous studies where human experts missed 1 in 4 breast cancer metastases in histopathology^27^, an issue that even increases substantially if humans work under time pressure^28^. Motivated by this, deep learning based approaches for cancer and metastasis detection recently gained substantial momentum for various imaging modalities, also beyond microscopy^29-32^.

Here we used an MDA-MB-231 cancer cell-based tumor model to train the algorithms. While training deep networks in general may require large training datasets to diversify their applications, the Unet-like architecture in the core of DeepMACT can be easily adopted to other cancer models^33-36^. In other words, DeepMACT learned to detect the characteristic shape and appearance of micrometastases against the background signal, and thus, is independent of the cancer cell model used. Therefore, it would require little effort to apply our algorithms to different types of tumor models. Also, adapting the algorithm to applications in which, for instance, shape and size differ substantially from MDA-MB-231 metastases, would not require training from scratch. Adjusting design parameters such as the size of subvolumes (see methods for details) allows the straightforward adaptation of the algorithm to new data with different SNR, metastasis sizes, or spatial resolution of the scan. Furthermore, building upon our pre-trained algorithms, which are freely available online, allows retraining the algorithm with substantially less training data.

To ensure high computational efficiency, our approach solves the three-dimensional task of detecting and segmenting the metastases by exploiting two-dimensional representations of the data. This is important because 2D max-intensity-projections increase SNR when there is little background noise owing to the high specificity of the labels in vDISCO clearing. 3D convolutions are exponentially more expensive in computing time than 2D convolutions, thus requiring more powerful computing resources and more data annotated in 3D to train the algorithm. Importantly, annotating projections in 2D is substantially easier and faster than slice-by-slice 3D annotations. In addition, the exponentially more efficient nature of our approach allows training the entire algorithm on a standard workstation with an ordinary GPU within a few hours; applying the trained algorithm to a new dataset takes in the order of 15 minutes, highlighting the scalability and cost-efficiency of our pipeline. Thus, this architecture is designed to enable widespread adaptation of our approach by minimizing data annotation and computing requirements while allowing for easy adaptation for other experimental setups (such as different imaging modalities or tumor models).

### DeepMACT detection of micrometastases and tumor-targeting drugs

Methods such as magnetic resonance imaging (MRI), computed tomography (CT), and bioluminescence imaging have been widely used to visualize cancer growth at the primary site and distal body regions^37-42^. While these methods provide crucial longitudinal information on the size of the primary tumor and large metastases, they typically can only resolve structures larger than 75 µm, hence they do not have the resolution to detect smaller micrometastases consisting of fewer cells.

Unbiased high-throughput mapping of tumor micrometastases at cellular resolution in entire rodent bodies can be a valuable tool to uncover the biology behind the dissemination of tumor cells. We show here that DeepMACT is an ideal tool for detecting and mapping cancer metastases in whole mouse bodies at the cellular level, allowing identification of the precise locations of single disseminated cancer cells. Complex analysis, e.g., of the size, location and density of micrometastases could be performed in a short time throughout the body, without dissecting any pre-defined region. In addition to detecting the micrometastases in the predicted organs such as the lungs and liver, we also identified numerous micrometastases throughout the torso. These may include cells that have metastasized to other organs, e.g. bones, peritoneum, bladder, intestines, as well as circulating tumor cells (or cells clusters) in the vasculature.

While precise assessment of antibody drug biodistribution is critical for evaluating its specificity and utility for tumor treatment, there have been no methods so far that can provide such information at the cellular level in the whole organism. Here, we applied DeepMACT to study not only the distribution of single tumor cells, but also of a therapeutic monoclonal antibody. We demonstrated that the “on” and “off” targeting of antibody drugs throughout the body can readily be assessed by DeepMACT. For example, we observed that not all micrometastases in the lungs were targeted by the anti-CA12 therapeutic antibody 6A10. Understanding why antibody-based therapeutics do not target all tumor cells would be important for developing more effective treatments. Towards this goal, we studied the potential mechanisms that could contribute to the lack of targeting. Vascular staining demonstrated that blood vessels were present in the immediate vicinity of all examined metastases in the lung, suggesting that insufficient vascularization is unlikely to be a common cause for the failure of antibody drug targeting. Interestingly, DeepMACT analysis found that micrometastases located in close proximity are more likely to be targeted. This suggests that the local microenvironment within metastatic niches plays an important role in determining the efficiency of antibody targeting, e.g. by altering antibody penetration, binding affinity and clearance. Furthermore, heterogeneity of antigen expression on the surface of tumor cells and internalization and degradation of antigen/antibody complexes might also affect therapeutic antibody targeting efficacy.

In conclusion, DeepMACT is a powerful technology combining unbiased whole mouse body imaging with automated analysis. It enables visualization, quantification, and analysis of tumor micrometastases and antibody-based therapies at single cell resolution in intact mice, with an accuracy equivalent to that of human experts, but speeding up the workflow by orders of magnitude compared to traditional methods. Because this technology is time- and cost-efficient, scalable, and easily adoptable, it can be used to study metastasis and optimize antibody-based drug targeting in diverse tumor models.

## ACKNOWLEDGMENTS

This work was supported by the Vascular Dementia Research Foundation, Synergy Excellence Cluster Munich (SyNergy), ERA-Net Neuron (01EW1501A to A.E.), the Helmholtz-Center for Environment Health (grants to R.Z.), and the German Federal Ministry of Education and Research via the Software Campus initiative (O.S.). Furthermore, NVIDIA supported this work with a Titan XP via the GPU Grant Program. We thank I. Jeremia for the luciferase/mCherry construct and the Monoclonal Antibody Core Facility at the Helmholtz Center Munich for providing the 6A10 antibody. We thank Dr. W. Ouyang for constructive comments on the manuscript. C.P. and R.C. are members of Graduate School of Systemic Neurosciences (GSN), Ludwig Maximilian University of Munich.

## AUTHOR CONTRIBUTIONS

C.P., A.P.D. and R.C. performed the tissue processing, clearing and imaging experiments. R.C. and C.P. developed the whole-body nanobody labeling protocol. M.T. stitched and assembled the whole mouse body scans. O.S. and J.D.B developed the deep learning architecture and performed the quantitative analyses. A.G. performed the image rendering and parts of the data analysis. O.S. developed annotation tools and the custom-made object detector, M.A.R. annotated the data. G.G.,B.N. and R.Z., performed tumor transplantation experiments and bioluminescence imaging. B.K.G. helped with data interpretation. B.M. provided guidance in developing the deep learning architecture and helped with data interpretation. A.E., C.P. and O.S. wrote the manuscript. All the authors edited the manuscript. A.E. initiated and led all aspects of the project.

## METHODS

### Spontaneous metastasis model and injection of therapeutic antibody

Female NSG (NOD/SCID/IL2 receptor *gamma* chain knockout) mice were obtained from Jackson Laboratory and housed at the animal facility of the Helmholtz Center Munich. All animal experiments were conducted according to institutional guidelines of the Ludwig Maximilian University of Munich and Helmholtz Center Munich after approval of the Ethical Review Board of the Government of Upper Bavaria (Regierung von Oberbayern, Munich, Germany). MDA-MB-231 breast cancer cells transduced with a lentivirus expressing mCherry and enhanced Firefly luciferase^43^ were counted, filtered through a 100 µm filter and resuspended in RPMI 1640 medium. 2×10^6^ cells per mouse were injected transdermally in a volume of 50 µl into the 4^th^ left mammary fat pad. Tumor growth was monitored by bioluminescence measurement (photons/second) of the whole body using an IVIS Lumina II Imaging System (Caliper Life Sciences) as described^19^. Briefly, mice were anesthetized with isoflurane, fixed in the imaging chamber and imaged 15 minutes after Luciferin injection (150 mg/kg; i.p.). Bioluminescence signal was quantified using the Living Image software 4.2 (Caliper). 9 weeks after tumor cell injections, mice were randomly assigned to different experimental procedures including injection of a human carbonic anhydrase (CA) XII-specific antibody (6A10)^24^, boosting of endogenous mCherry fluorescence, immunolabeling and clearing, as described below. 48 hours before perfusion mice were injected into the tail vein with 20 µg of 6A10 antibody conjugated with Alexa-568.

### Perfusion and tissue processing

Mice were deeply anesthetized using a combination of midazolam, medetomidine and fentanyl (MMF) (1ml/100g of body mass for mice; i.p.) before intracardial perfusion with heparinized 0.01 M PBS (10 U/ml of Heparin, Ratiopharm; 100-125 mmHg pressure using a Leica Perfusion One system) for 5-10 minutes at room temperature until the blood was washed out, followed by 4% paraformaldehyde (PFA) in 0.01 M PBS (pH 7.4) (Morphisto, 11762.01000) for 10-20 minutes. The skin was carefully removed and the bodies were postfixed in 4% PFA for 1 day at 4 °C and transferred to 0.01 M PBS. The vDISCO pipeline was started immediately or whole mouse bodies were stored in PBS at 4 °C for up to 4 weeks or in PBS containing 0.05% sodium azide (Sigma, 71290) for up to 6 months.

### uDISCO whole-body clearing

The uDISCO protocol to clear whole body of mice was already described in details in ref (Pan et al., 2016). In brief, a transcardial-circulatory system was established involving a peristaltic pump (ISMATEC, REGLO Digital MS-4/8 ISM 834; reference tubing, SC0266). Two channels from the pump were set for the circulation through the heart into the vasculature: the first channel pumped the clearing solution into the mouse body and the second channel collected the solution exiting the mouse body and recirculated the solution back to the original bottle. For the outflow tubing of the first channel, which injected the solution into the heart, the tip of a syringe (cut from a 1 ml syringe-Braun, 9166017V) was used to connect the perfusion needle (Leica, 39471024) to the tubing. Meanwhile, the inflow tubing of the second channel, which recirculated the clearing solutions, was fixed to the glass chamber containing the mouse body. The amount of solutions for circulation depended on the capacity of the clearing glass chamber. For example, if the maximum volume of glass chamber is 400 ml, 300 ml of volume of solution was used for circulation.

All clearing steps were performed in a fume hood. Firstly, the mouse body was put in a glass chamber and the perfusion needle was inserted into the heart through the same hole that was used for PFA perfusion. Then, after covering the chamber with aluminum foil the transcardial circulation was started with a pressure of 230 mmHg (60 rpm on the ISMATEC pump). The mouse bodies were perfused for 6 hours with the following gradient of *tert*-butanol: 30 Vol%, 50 Vol%, 70 Vol%, 90 Vol% (in distilled water),100 Vol% twice, and finally with the refractive index matching solution BABB-D4 containing 4 parts BABB (benzyl alcohol + benzyl benzoate 1:2, Sigma, 24122 and W213802), 1 part diphenyl ether (DPE) (Alfa Aesar, A15791) and 0.4% Vol vitamin E (DL-alpha-tocopherol, Alfa Aesar, A17039), for at least 6 hours until achieving transparency of the bodies. As the melting point of *tert*-butanol is between 23 to 26 °C, a heating mat set at 35-40 °C was used for the two rounds of 100% *tert*-butanol circulation to prevent the solution from solidifying.

### vDISCO whole-body immunostaining and clearing

The detailed protocol of vDISCO is described in ref (Cai et al, 2018). The following nanobodies and dyes were used for whole body immunostaining: Atto647N conjugated anti-RFP/mCherry nanobooster (Chromotek, rba647n-100), Atto594 conjugated anti-RFP/mCherry nanobooster (Chromotek, rba594-100), Hoechst 33342 (Thermo Fisher Scientific, 21492H), Propidium iodide (PI, Sigma, P4864).

After PBS perfusion and PFA fixation, the animals were placed into a 300-ml glass chamber and the same transcardial-circulatory system with a peristaltic pump was established to perfuse the mice during decolorization and immunostaining steps. The animals were firstly perfused with decolorization solution for 2 days at room temperature to remove remaining heme and blood before immunostaining. The decolorization solution which is a 1:3 dilution of CUBIC reagent 1 (Susaki et al., 2014) in 0.01 M PBS was refreshed every 12 hours. CUBIC reagent 1 was prepared as a mixture of 25 wt% N,N,N,N’-tetrakis (2-hydroxypropyl) ethylenediamine (Sigma-Aldrich, 122262), 25 wt% urea (Carl Roth, 3941.3) and15 wt% Triton X-100 in 0.1 M PBS, as described in the original publication. Before the immunostaining step, additional 0,22 µm syringe filters (Sartorius 16532) were attached to the tubing to prevent the potential accumulation of nanobody aggregates and ∼230 mmHg high pressure pumping was maintained through the entire labeling process. Subsequently the animals were perfused for 5-6 days at room temperature with 300 ml of immunostaining solution containing 0.5% Triton X-100, 1.5% goat serum (Gibco, 16210072), 0.5 mM of Methyl-beta-cyclodextrin (Sigma, 332615), 0.2% trans-1-Acetyl-4-hydroxy-L-proline (Sigma, 441562), 0.05% sodium azide (Sigma, 71290), 25 µL of nano-booster (stock concentration 0.5 – 1 mg/ ml), 10 µg/ml Hoechst and/or 350 µL of propidium iodide (stock concentration 1mg/ml) in 0.01 M PBS. Then the mice were perfused with washing solution (1.5% goat serum, 0.5% Triton X-100, 0.05% of sodium azide in 0.01 M PBS) for 12 hours twice at room temperature and at the end with 0.01 M PBS for 12 hours twice at room temperature.

After the whole body immunolabeling, the whole mouse bodies were passively cleared using 3DISCO. In short, mice were cleared at room temperature inside a glass chamber with gentle shaking under a fume hood. For dehydration, mice bodies were incubated in 250 ml of the gradient tetrahydrofuran THF (Sigma, 186562) in distilled water (6-12 hours for each step): 50 Vol% THF, 70 Vol% THF, 80 Vol% THF, 100 Vol% THF and again 100 Vol% THF; then mice were incubated for 1 hour in dichloromethane (Sigma, 270997), and finally in BABB. During all clearing steps, the glass chamber was sealed with parafilm and covered by aluminum foil. Further details on the vDISCO protocols are available at http://discotechnologies.org/vDISCO/.

### Lectin vasculature labeling in lungs tissue sections

The whole bodies of mice were perfused and collected as described above. After checking with epifluorescence stereomicroscopy (Zeiss AxioZoom EMS3/SyCoP3), the lung lobes with multiple metastases were dissected and sliced into 20 µm thick tissue sections by using a cryostat (Leica, CM3050S). The lung sections were washes 2 times with 0.01 M PBS and then incubated in Alexa 488 conjugated Lectin (4 µg/ml, invitrogen, W11261) at 4 °C overnight. The sections were then stained with Hoechst 33342 (10 µg/ml, Thermo Fisher Scientific, 21492H) for 5 minutes at room temperature to visualize the nucleus. After washing 2 times with PBS, the slides were mounted with fluorescent mounting medium (Dako, 10097416) and were ready for confocal microscopy.

### mCherry nanoboosting in lung tissue sections

20 µm thick lung tissue sections were washed with 0.01 M PBS 2 times before starting the boosting process. One hour incubation in blocking solution containing 1% Bovine Serum Albumin (Sigma, A7906), 2% goat serum (Gibco, 16210-072), 0.1% Triton X-100 and 0.05% Tween 20 (Bio-Rad, 161-0781) in PBS, was performed at room temperature. Then the staining solution was prepared in 1% Bovine Serum Albumin and 0.5% Triton X-100 in PBS. Atto647N conjugated anti-RFP/mCherry nanobooster was diluted 1:500 in the staining solution and the lungs sections were incubated overnight at 4°C. After the nanoboosting, the lungs sections were washed 3 times with PBS for 5 minutes with gentle shaking. After nuclear staining by Hoechst 33342 (10 µg/ml) and post wash with PBS as described before, the slides were mounted with fluorescence mounting medium and were ready for confocal microscopy.

### Epifluorescence stereomicroscopy imaging

Cleared whole mouse bodies were fixed in the original clearing chamber and were imaged with ZeissAxioZoomEMS3/SyCoP3fluorescence stereomicroscope using a 1x long working distance air objective lens (Plan Z 1x, 0.25 NA, Working distance (WD) = 56 mm). The magnification was set as 7x and imaging areas were selected manually to cover the entire mouse bodies. The images were taken with GFP, RFP and Cy5 filters and files were exported as RGB images.

### Light-sheet microscopy imaging

Single plane illumination (light-sheet) image stacks were acquired using an Ultramicroscope II (LaVision BioTec), allowing an axial resolution of 4 μm. For low magnification whole-body screening of tumor and antibody signals we used a 1x Olympus air objective (Olympus MV PLAPO 1x/0.25 NA [WD = 65mm]) coupled to an Olympus MVX10 zoom body, which provides zoom-out and -in ranging from 0.63x up to 6.3x. Using 1x objective, we imaged a field of view of 2 x 2.5 cm, covering the entire width of the mouse body. Tile scans with 60% overlap along the longitudinal y-axis of the mouse body were obtained from ventral and dorsal surfaces up to 13 mm in depth, covering the entire volume of the body using a z-step of 10 µm. Exposure time was 150 ms, laser power was 3 to 4 mW (70% to 95% of the power level) and the light-sheet width was kept at maximum. After low magnification imaging of the whole body, individual organs (including lungs, liver, kidneys, brain, spleen, intestines and bones) were dissected and imaged individually using high magnification objectives (Olympus XLFLUOR 4x corrected/0.28 NA [WD = 10 mm] and Zeiss 20x Clr Plan-Neofluar/0.1 NA [WD 4 = mm]) coupled to an Olympus revolving zoom body unit (U-TVCAC) kept at 1x. High magnification tile scans were acquired using 20% overlap and the light-sheet width was reduced to obtain maximum illumination in the field of view keeping the same NA. For the data used for the comparison of signal profile plots of lung metastases taken in red and far-red channels and for the analysis of endogenous fluorescence signal depletion after the uDISCO protocol, we used the same MVX10 zoom body, coupled this time with a 2x objective (Olympus MVPLAPO2XC/0.5 NA [WD = 6mm]) at zoom body magnification 6.3x and 2.5x respectively.

### Confocal microscopy imaging

For imaging the thick cleared specimens such as dissected tissues, pieces of organs or whole organs were placed on 35 mm glass bottom petri dishes (MatTek, P35G-0-14-C), then the samples were covered with one or two drops of the refractive index matching solution such as BABB or BABB-D4. Sealing of this mounting chamber was not necessary. The samples were imaged with an inverted laser-scanning confocal microscopy system (Zeiss, LSM 880) using a 40x oil immersion lens (Zeiss, EC Plan-Neofluar 40x/1.30 Oil DIC M27) and a 25x water immersion long-working distance objective lens (Leica, NA 0.95, WD = 2.5mm), the latter one was mounted on a custom mounting thread. The z-step size was 1-2.50 μm. For imaging the lung tissue sections with lectin staining and with nanoboosters, the slides were imaged with the same inverted laser-scanning confocal microscopy system (Zeiss, LSM 880) using a 40x oil immersion lens (Zeiss, EC Plan-Neofluar 40x/1.30 Oil DIC M27). The z-step size was 2 µm.

### Reconstructions of whole-mouse body scans

#### Epifluorescence (2D montage of whole mouse)

The collected epifluorescence images were stitched semi-automatically using Adobe Photoshop photomerge function (File\automate\photomerge). The different channels were stitched separately and merged in Adobe Photoshop to generate the composite images.

#### Light-sheet microscopy (3D montage of whole mouse)

We acquired light-sheet microscope stacks using ImSpector (LaVision BioTec GmbH) as 16-bit grayscale TIFF images for each channel separately. The stacks were first aligned and fused together with Vision4D (Arivis AG). Further image processing was done mostly in Fiji (ImageJ2): first, the autofluorescence channel (imaged in 488 excitation) was equalized for a general outline of the mouse body. The organs were segmented manually by defining the regions of interests (ROIs). Data visualization was done with Amira (FEI Visualization Sciences Group), Imaris (Bitplane AG), Vision4D in both volumetric and maximum intensity projection color mapping.

### General data processing

All data processing after image volume reconstruction was performed in Python using custom scripts based on publicly available standard packages comprising SciPy^44^, Seaborn^45^, and Pandas^46^. Deep Learning models were build using the PyTorch framework^47^. Since a single whole body scan is in the order of several terabytes due to its high resolution (the data used for training had a voxel size of (10µm)^3^), the volume was divided into 1176 subvolumes of (350px)^3^ (or (3.5mm)^3^) to enable efficient processing. Subvolumes were overlapping by 50px to ensure any given metastasis is fully captured by at least one subvolume to avoid artefacts of divided metastases at subvolume interfaces. Please note that the size and overlap of subvolumes are design choices that allow easy adaptation to different data sets, e.g. with different SNR, metastasis sizes, or spatial resolution of the scan. Final analyses were conducted on the re-assembled full volume whereby reconcatenation ruled out any double-counting at previously overlapping subvolumes.

### Data annotation by human experts

To provide ground truth in the form of a commonly agreed upon reference annotation for training, as well as for evaluation of the algorithms developed, full body scans of two mice were manually annotated by a group of human experts. This manual process was augmented with a set of tools to reduce the total workload from an estimated total duration of several months down to 150 person-hours net annotation time.

#### Automatic pre-annotation with custom-made filter-based detector

To avoid starting from scratch to annotate two volumes of several thousand z-slices, an automatic detection and segmentation method was applied to provide a basis for manual correction. Due to the insufficient performance of established methods (in this case: the 3D Object Detector for ImageJ^48^, we developed a custom-made filter based detector tailored to the specifics of this dataset. In brief, we handcrafted a spatial filter kernel optimized to detect the most common metastases and applied it with 3D convolutions to the dataset; subsequent binarization and connected-component analysis yielded *seed points* collocated with metastases. This allowed for further analyses of the immediate local neighborhood of these candidate regions; a local 3D segmentation was derived by selective region growing around these seed points based on the local signal intensity distribution up to a mean *foreground signal* limited to 4 standard deviations above the mean signal in the local surrounding. Finally, obvious false positives were filtered out. Together, this approach generated a first proposal for the data annotation that at least captured the most obvious metastases while producing an acceptable rate of false positives. As shown in the results section, the quality of this proposal was about twice as good as compared to the 3D Object Detector in ImageJ (35% instead of 18% in F1-score). Importantly, further fine-tuning of filters and parameters and any additional automated pre- or post-processing did not improve the results, indicating that a F1-score of 35% may be close to the performance limit of such approaches with fixed filter kernels and fixed decision rules for such kind of data.

#### Manual annotation correction by human experts

This first proposal served as a basis for human annotation. In general, three kinds of manual correction were needed to derive a good annotation: removal of false positives, addition of false negatives (previously missed metastases) and adjustment of the 3D segmentation of each metastasis. To avoid the need to perform this task individually for each of the 350 layers of a (350px)^3^ data subvolume, a custom tool with an interactive graphical user interface was developed. Based on maximum intensity projections along each dimension, the tool allowed to review, adjust, add, and remove each potential metastasis in the subvolume with a few mouse clicks, drastically speeding up the annotation process from hours to minutes per subvolume. Different perspectives (X, Y, Z) and viewing modes (e.g., projections, orthogonal slices, adjusted contrasts, 3D renderings) for each individual metastasis allowed the annotator to take maximally informed decisions even in less obvious cases.

#### Refinement of annotation to commonly agreed upon ground truth

A small fraction (3%) of the entire data set was labeled several times by the annotators without their awareness to assess human labeling consistency. Since the difference in annotation for a given subvolume between two trials of a single annotator was about as big as between two independent annotators and quite substantial (the agreement between two trials of the same annotator or between annotators only reached an F1-score of 80-85%) we decided to invest additional time to refine the entire data set. First, all experts (3 graduate students with extensive experience in the field of imaging and tumor biology) jointly discussed examples of annotation differences to build a common understanding. Annotations of subvolumes with the biggest discrepancies were again reviewed and refined. Furthermore, this analysis revealed that the most prevalent source of annotation error was overlooked metastases (false negatives). Here, around 29% of metastases were missed in the human annotation, in line with previous studies^27,28^. To effectively identify all missed metastases in the entire data set, our deep learning algorithm (see next section) was trained on the status quo of the annotations and applied to the data set with high sensitivity. This yielded a long list of potential candidates. With the help of another custom-built, interactive graphical user interface, all potential candidates were manually reviewed by the annotators and either discarded or manually adjusted and added to the segmentation. A small set of potential metastases, for which human annotators could not take a conclusive decision even after joint discussion, was recorded separately, but not added to the segmentation. These laborious steps ensured the generation of a high-quality *ground truth* for training the algorithm and, importantly, for evaluating its performance in comparison to a single human annotator. Here, this selectively iterative approach of refining annotations based on the input of several human experts was chosen due to the substantial amount of manual work involved with reviewing our high-resolution scans. Since a full review of one person takes about a month of full-time work, repeating this process several times would be desirable but too costly. In applications where several, independent full annotations are available, advanced mathematical frameworks for refining decisions from different experts to a single decision can be applied in order to avoid a bias towards individual decisions^49,50^.

### Deep learning algorithm for metastasis detection and segmentation

#### Implementation details of the model architecture

Inspired by the established U-net architecture^21^, we designed a deep learning approach that is depicted in **Figures 3A** and briefly described in the results section. The architecture of the CNN at its core (**Figure 3C**) is characterized by an encoding downward path and a decoding upward path comprising a total of 7 levels, in which each level also has a lateral skip-connection that bypasses the deeper levels and feeds the output of the encoding unit directly to the corresponding decoding unit. Each encoding unit increases the number of feature channels per pixel with the help of two kernel-based convolutions (kernel size: 3; padding: 1; dilation: 1; stride: 1) followed by batch normalization and a rectifying linear unit (ReLU). While the first convolutional step increases the number of feature channels, this number stays constant for the second convolutional step. Before being passed on to the next encoding stage, the spatial resolution is halved using max-pooling (kernel size: 2, stride: 2). Decoding units take two inputs: the output from the previous layer is spatially upsampled by a factor of two (bilinearly) and concatenated along the feature dimension with the output of the corresponding encoding unit, bypassing the deeper levels. A first convolutional step (same parameters as before) decreases the number of feature channels, which is again kept constant in the two subsequent convolutions. The 24-feature channel output of the last decoder is mapped to logits in the 2D space with a convolutional step without padding, batch normalization, or a rectifying linear unit.

#### Training, validation and test sets

Following established standards, model training and evaluation was based on k-fold cross-validation (k=5). Thus, the data set was split into mutually exclusive sets for training and validation (80%) and for testing (20%). This process was repeated k times, yielding a total of 5 mutually exclusive test sets that are collectively exhaustive. The network weights and all design choices and hyperparameters (such as batch size, learning rate, etc) were optimized solely with the training and validation set to avoid overfitting on the specifics of the test set. The data set was confined to subvolumes within the torso of the mouse body as subvolumes containing near-zero values outside the body contain no useful information to train or test on. In contrast to all metastases in the entire body, the tumor tissues of the primary tumor and the auxiliary lymph node are several orders of magnitude larger (i.e., they follow very different statistics than all micrometastases) and were thus excluded. The signal from one subvolume was corrupted by a dirt particle and thus also excluded. In total, these exclusions made up less than 1% of the total scan volume. The split between the three subsets (training, validation, testing) of the data was done on a subvolume level (from which the three projections are created afterwards) to avoid information leak between different projections from the same subvolume.

#### Training and validation procedure

The model training was conducted in two steps. First, a large number of models spanning a broad set of different hyperparameters were trained for 10 epochs using another (nested) k-fold cross-validation (k=5) within the training and validation set. Second, the model with the best-performing set of hyperparameters (presented here) was trained for the remaining epochs. Thus, any hyperparameter choice was made without looking at the performance on the test set. The model was trained for 40 epochs of the entire training data set, using random vertical and horizontal flips of the data to augment its variance (further training epochs did not improve the predictive power). We used a batch size *B* of 4 but found that other batch sizes work similarly well. Each input was normalized by its local subvolume peak value, which was found to work better than normalization to the global volume peak value or non-linear normalizations. To calculate the gradients for network weight optimization (i.e., to train the model), we used weighted binary cross entropy as a loss function for a given prediction Ŷcompared to the ground truth *Y*, giving more weight *w* for foreground (*FG*) pixels *p* versus background pixels (*BG*) to account for the *class imbalance* (i.e., that metastases are very sparsely distributed in space):

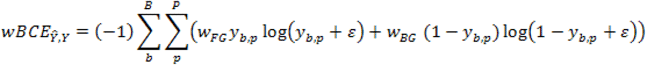

A small numerical offset ^ε^ *= 10*^*-4*^ was applied for numerical stability. We found equal weights or a slightly stronger bias to foreground to work almost equally well (here, we used *w*_*FG*_*=2* and *w*_*BG*_*=0.5*), larger biases had negative effects. Additionally, we allowed the network to optimize the share of training data that contains at least some foreground by ignoring parts of training data in which no foreground is present. A share of 90% training data with at least some foreground optimized the performance on the validation set and was thus chosen. The network was trained using the Adam optimizer^51^; the initial learning rate was set to 10^-4^ and was gradually decreased by a factor of 10 to a minimum of 10^-7^ every time the loss function reached a plateau for more than 2 epochs. A single training run over 40 epochs takes only around 20-30 minutes on a normal workstation equipped with a NVIDIA Titan Xp GPU.

#### Testing and inference mode

As mentioned before, we applied k-fold cross-validation. Thus, in each of the *k=5* folds the model was tested on data that was not seen by the model during training and validation. Together, all 5 test sets span the entire data set. As depicted in **Figure 3A**, the trained algorithm was used to generate probability masks for each of the three projection perspectives (*P*_*XY*_, *P*_*YZ*_, *P*_*ZX*_), in which the pixel value indicates the network’s confidence t hat t his pixel is part of a metastasis in the given sub-volume *s*. Building the outer product of the three probability masks allows to recombine the three somewhat independent judgements of the network in 3D space:

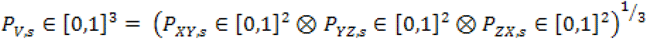

This 3D recombination *P*_*V*_ of the 2D probability maps yields a final predicted segmentation mask after binarization. By default, the confidence threshold was set to 50%; however, changing this parameter allows to manually adjust the trade-off between sensitivity and specificity, if desired (also see **Figure S4**). Please note that the F1-score for evaluation is not affected by this trade-off (i.e., a better detection rate at the cost of a higher false positive rate would not artificially increase the F1-score and vice versa). Subsequent connected-component analysis converts the output to an explicit list and segmentation of predicted metastases in 3D space.

#### Performance evaluation

The same performance evaluation procedure was used for the comparison shown in **Figure 3D**, including the performance of a single human annotator. A standard test for detection tasks, the F1 score quantifies the accuracy of a model by combining precision (share of true positives among all positive predictions, including false positives) and recall (share of predicted positives among the sum of the true positive and false negative predictions). It is mathematically equivalent to the Sørensen–Dice coefficient (“Dice score”), which is the commonly used name for pixel-wise image segmentation problems. The F1 score is given as:

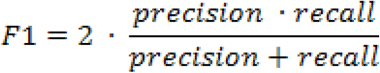

For all comparisons of detection and segmentation performance, the ground truth (refined by several human experts as described above) is used as a reference. We quantified the performance of the proposed deep learning algorithm based on its prediction of the test set. For a comparison, the segmentations as provided by established tools like the 3D Object Detector in ImageJ, our custom-made detector as described above, as well as the annotation as provided by a single human annotator (before joint refinement) were quantified in the same manner. Overlapping segmentations for metastases were counted as true positive predictions, non-overlapping predictions as false positives and metastases not detected by the prediction as false negatives. Predictions corresponding to the small set of cases unclear to the group of human experts (see above) were neither counted as true positive predictions nor as false negatives, i.e. they neither increased nor decreased the performance evaluation. All performance evaluations were conducted on the entirety of the test set as a whole. To quantify the inherent variance, the distribution of performance results was estimated with n=1000 resampled test sets (of same size) using the bootstrapping approach.

The DeepMACT algorithms can be downloaded at: http://discotechnologies.org/DeepMACT.

### Quantifications and statistical analysis

#### Organ registration

For the full body light-sheet scans (e.g., **Figure 4C,F**) the outlines of selected organs of interest (all lung lobes, both kidneys, liver) were manually segmented as multi-point polygons in a stack of slices in 3D using Fiji. For each metastasis detected by our deep learning architecture we assessed whether its center of mass falls into the 3D segmentation of one of those organs using a custom Python script. Any metastasis not registered to one of these organs is referred to as located in “the rest of the torso” in this manuscript. The 3D segmentation of the lungs was also used to compute the overall lung volume to assess the tumor density in **Figure 4J**, which we quantified as the share of the sum of the volume of all metastases registered to an organ of the entire organ volume.

#### Metastasis characterization

The output of our deep learning architecture is a binary segmentation volume for all metastases. We developed a custom-made, highly efficient implementation of a connected component analysis (available online) to derive an explicit list of metastases fully characterized in 3D. Based on each metastasis 3D shape and voxel-based volume *V*, we computed its average diameter as

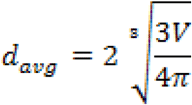

To quantify the number of cells in each metastasis, we measured the size of several isolated single cells (∼1700 µm^3^) and estimated the total numbers in metastases based on volumetric interpolation. We confirmed the accuracy of the estimations by the number of nuclei (PI or Hoechst labeled) in selected micrometastases. The distance of each metastasis *i* to its nearest neighboring metastasis was measured in 3D space as the Euclidian distance between their center of masses *CoM*:

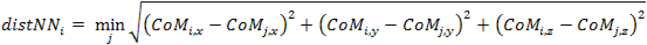

#### Drug targeting analysis

We assessed the 6A10 antibody targeting of a given metastasis by analyzing the distribution of the fluorescent signal strength within the 3D segmentation of each metastasis (*ξ*_*m*_) versus the distribution in its local surrounding (250 µm around the metastasis) *ξ*_*s*_. For each signal distribution, the number of voxels within the metastasis segmentation *n*_*m*_or in its local surrounding *n*_*s*_can be seen as the number of observations of the underlying true (but unknown) distributions. The degree of targeting was estimated by quantifying the ratio of mean signal strength within the segmentation to the mean signal strength in its surrounding (e.g., in **Figure 5K**). We refer to this as antibody signal ratio. A ratio larger than 1 means that the antibody signal strength within then 3D segmentation of the metastasis is higher than around it (see dashed line in **Figure 5K**). Whether or not a metastasis was deemed “targeted” was assessed with a version of the *t*-test to determine whether mean of the observed signal distribution in the metastasis *ξ*_*m*_was significantly at least *Δ=50% (ratio of 1.5)* above the mean of the observed signal distribution in the local surrounding *ξ*_*s*_. Importantly, a *t*-test is valid for the signals despite their highly non-normal underlying distribution as the number of observations far exceeds the requirements of the central limit theorem (i.e., while the signals are not normally distributed, the estimation of their means is normally distributed due the high number of observations). This was verified manually. However, due to a typically much larger number of observations in the local surrounding *ξ*_*s*_ than for the metastasis itself *ξ*_*m*_, the statistical test was not performed with a *Student’s t-test* but with the *Welch’s t-test* that corrects the *degrees of freedom* for an unequal number of observations for both distributions:

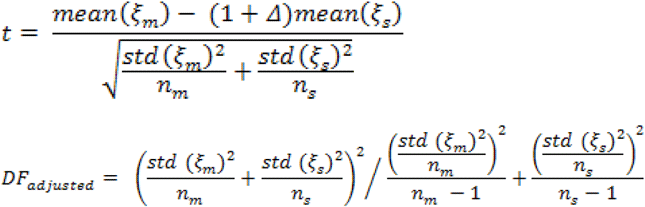

#### Analysis of fluorescence signal profiles from light-sheet images

We considered the fluorescence signal profiles from each channel: excitation 470 nm, 561 nm and 647 nm. These profiles were plotted in the same z-stack and normalized as percentage over the maximum peak. To compare the reduction of the background and the improvement of the signal over background ratio (SBR) in far-red and near far-red channels, we analyzed lung metastases expressing mCherry imaged with excitation 545/561 nm, lung metastases labeled with anti-mCherry nanobody conjugated with Atto594 imaged with excitation 590 nm, and lung metastases labeled with anti-mCherry nanobody conjugated with Atto647N imaged with excitation 640 nm (n=9 tumors per each experimental group which consisted of 3 animals per each imaging modality). The signal profile was measured from a defined straight line covering the tumors and surrounding tissue background and all the values of the plot from a representative animal per each experimental group were shown in a representative line chart (**Figure S1D**). Finally, the normalized plots represented in **Figure S1E** were calculated by normalizing the plots of lung metastases obtained as described above over the average signal intensity of the respective surrounding background.

#### Quantification of metastasis diameter and distance between metastases and vessels

Metastasis diameters were verified manually. For quantifying the distance between metastases and vessels, ten points on the border of each metastasis were randomly selected and the shortest distance from these points to the closest vessel wall were measured. The presented distance between each metastasis and nearest vessels was quantified by averaging these ten measurements. In Figure 6B, 50 metastases were quantified to generate the distribution map and each bullet point represent one single metastasis.

## MOVIE LEGENDS

### Movie S1

3D reconstruction of an intact mouse torso scanned by light-sheet microscopy. The left side shows the segmented view, in which the torso is represented in gray, the lungs in yellow and the heart in green. The right side shows the original 2D data from the light-sheet microscopy. Various macro-and micro-metastases (magenta) are visualized in high contrast over the background.

### Movie S2

3D visualization of some micrometastases in the lungs showing the single cell resolution obtained by light-sheet microscopy. The tumors are shown in magenta and PI nuclear labeling in green.

### Movie S3

3D animation of whole mouse body scanned by light-sheet microscopy at the cellular level. The outline of the mouse (scan of the unlabeled autofluorescence channel at 488 nm excitation) shown in gray, some segmented organs (lungs, kidney, liver and brain) in blue, the tumors in magenta, and the tumor & therapeutic antibody 6A10 co-localization in white. The first part of the animation shows tumor macro- and micrometastasis throughout the body, and the second part shows the biodistribution of the therapeutic antibody 6A10 along with tumor macro- and micrometastases.

### Movie S4

3D animation of the lungs demonstrating the details of the therapeutic antibody binding. Micrometastases are shown in magenta and the co-localization of micrometastases and therapeutic antibody in white. While most of the micrometastases are targeted by the therapeutic antibody (white), a fraction was not (the ones remaining magenta throughout the movie).

---

**Figure S1; related to Figure 2.**
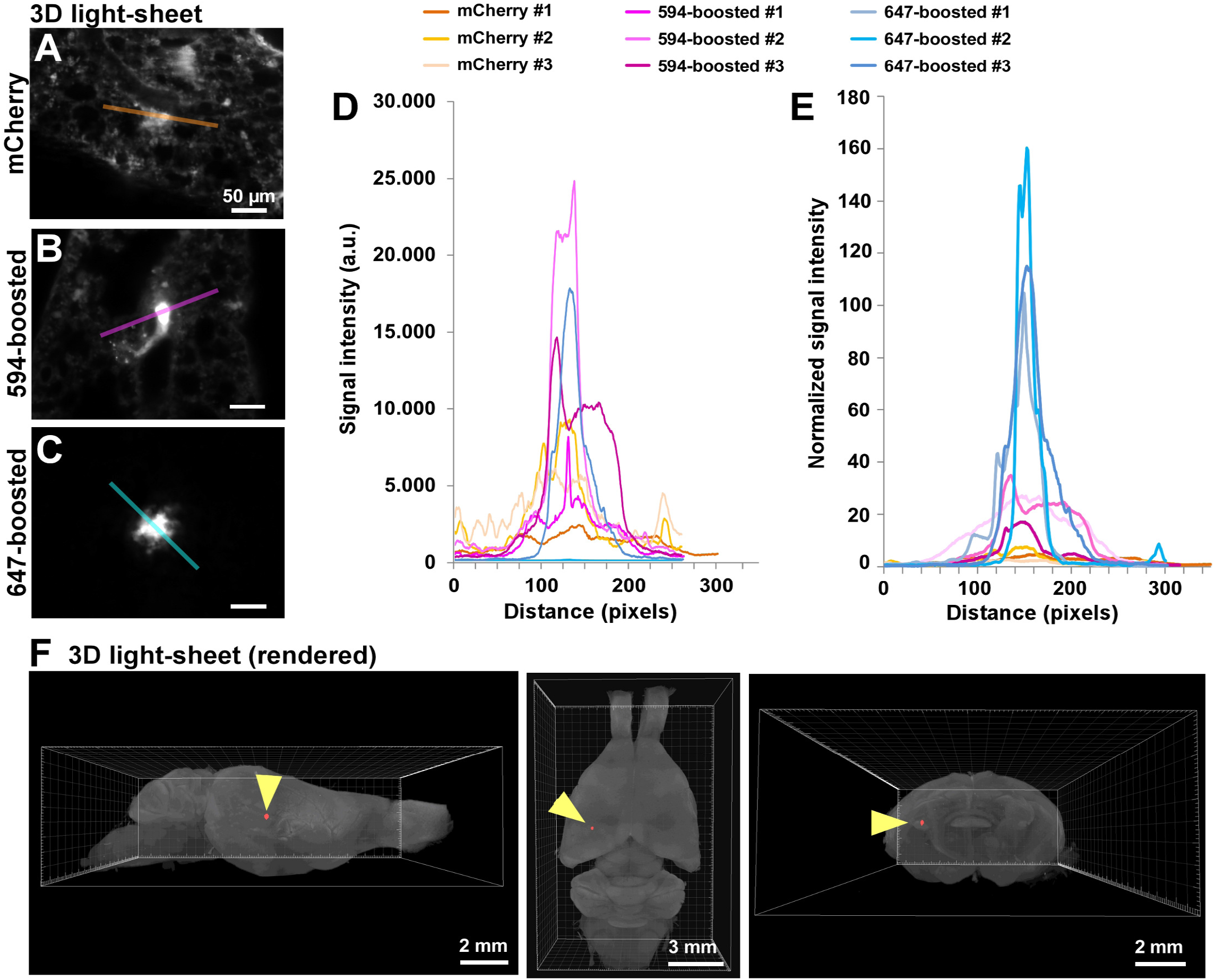
vDISCO nanoboosting of the fluorescent signal of cancer cells (A-C) Representative light-sheet images of mCherry expressing tumor metastases in the lungs of cleared mice that were not nanobody-boosted **(A)**, metastases boosted with an anti-mCherry nanobody conjugated to Atto594 **(B)** or an anti-mCherry nanobody conjugated to Atto647N **(C). (D)** Plots of signal intensity profiles along the yellow lines in panels A-C: non-boosted mCherry (orange), mCherry boosted with Atto594 (magenta) or mCherry boosted with Atto647N (cyan) (n=3 representative meta stases). **(E)** Intensity profiles of the fluorescence signal in (D) normalized over the background. **(F)** Example of deep-tissue imaging of Atto647N-boosted tumor metastases in the brain in the far-red spectrum of transparent mice after vDISCO. A tumor micrometastasis, which is located several millimeters deen in the brain tissue. can be readily detected (yellow arrowhead.

**Figure S2; related to Figure 2.**
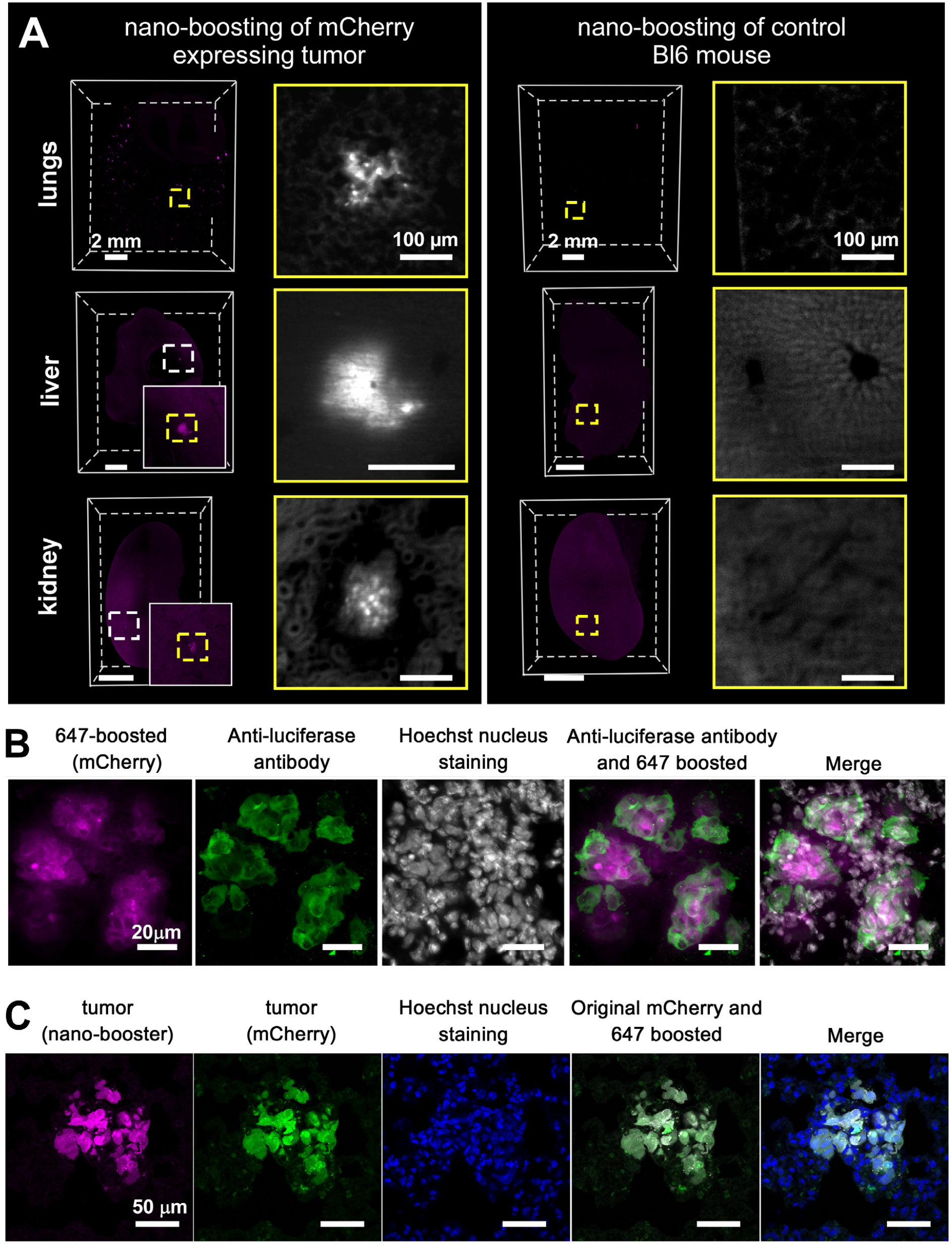
Confirmation of specificity of nano-booster staining ¡n mice bearing mCherry expressing tumors. **(A)** Comparison between an animal bearing an mCherry expressing tumor and a C57BLJ6N control animal, which were both boosted with an anti-mCherry nanobody conjugated to Atto647N (magenta) and imaged by light-sheet microscopy. No signal is detected in organs from the C57BL/6N control. Note that the background is enhanced to demonstrate the absence of signal in the high-magnification images. **(B)** Confocal images of metastatic lung tissue immunolabeled with an anti-firefly luciferase antibody (green) after rehydration of the cleared tissue; Atto647N-boosted cancer cells and cell nuclei are shown in magenta and grey,respectively. **(C)** Confocal images of a metastasis in the lung of an animal labeled with an anti-mCherry nano-booster conjugated to Atto647N. The nano-booster is shown in magenta,mCherry is shown in green and cell nuclei are shown in blue indicating that nano-boosting specifically detects mCherry.

**Figure S3; related to Figure 2.**
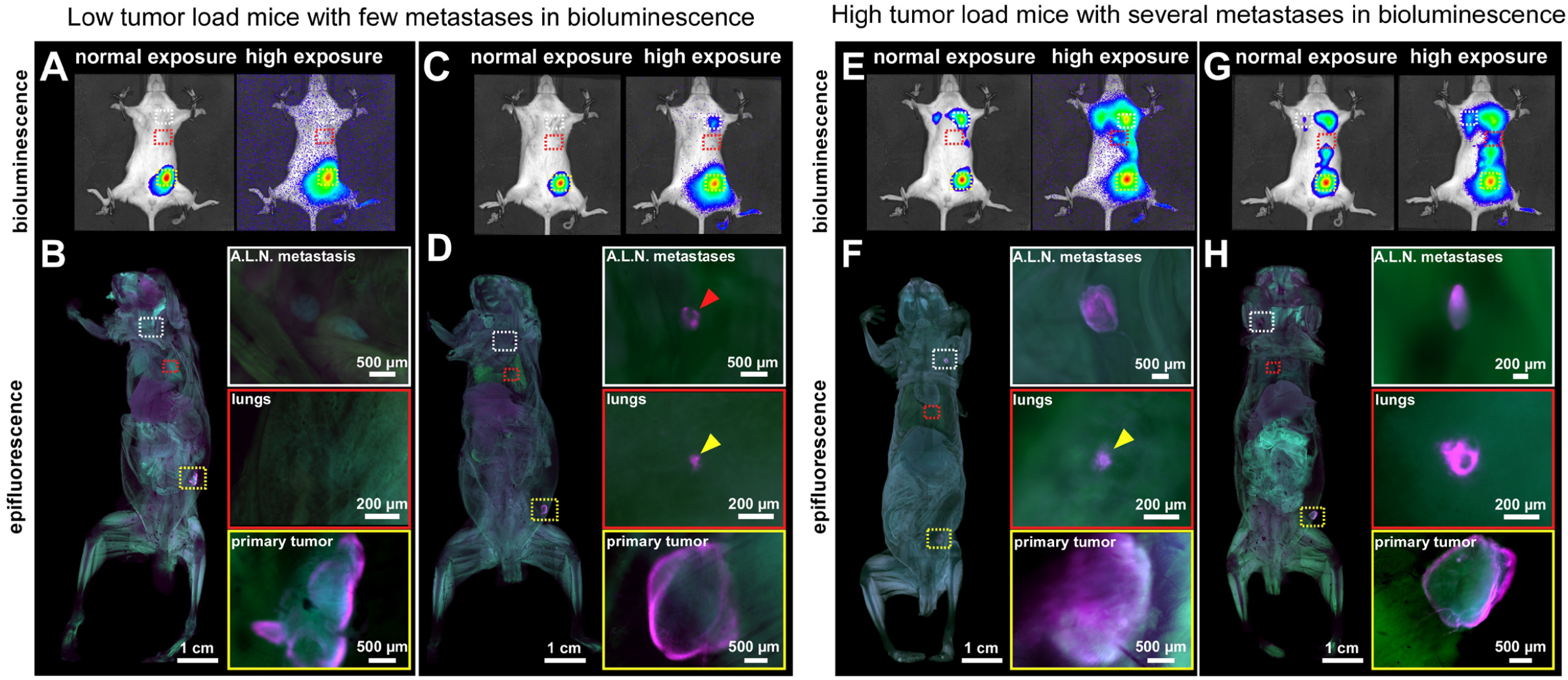
Examples of tumor metastasis detection in mice using bioluminescence imaging versus vDISCO and epifluorescence microscopy. **(A-D)** We found that bioluminescence imaging with normal exposure is not sufficiently sensitive to detect all the metastases in low tumor load mice. For example, the mice in **(A,B)** and **(C,D)** had very similar bioluminescence images with normal exposure. Applying vDISCO to these mice, we found no tumor metastases in one case **(A,B)** and a large metastasis (red arrowhead) in axillary lymph nodes (A.L.N. metastasis) **(C,D)** using a fluorescence stereo microscope. Although the signal from the primary tumor is strong in both normal and high exposure bioluminescence images (**A,C,.** yellow box), metastases in lungs (**A.C**, red boxes) are not visible, but are detected by epifluorescence imaging (D, yellow arrowhead). In epifluorescence images, the tumors (A647 labeled) are shown in magenta and the background, scanned in 488 nm, is shown in green. **(E-H)** In mice with high tumor load, a bulk heat map of metastatic distribution can be obtained by in the bioluminescence imaging, without detailed shape and size information. In contrast, vDISCO resolved single micrometastases in whole mouse bodies even with a fluorescence stereo microscope. Especially in the lungs,even micrometastases with a diameter smaller than 100 pm could be resolved in intact mice (**F**, yellow arrowhead).

**Figure S4; related to Figure 3.**
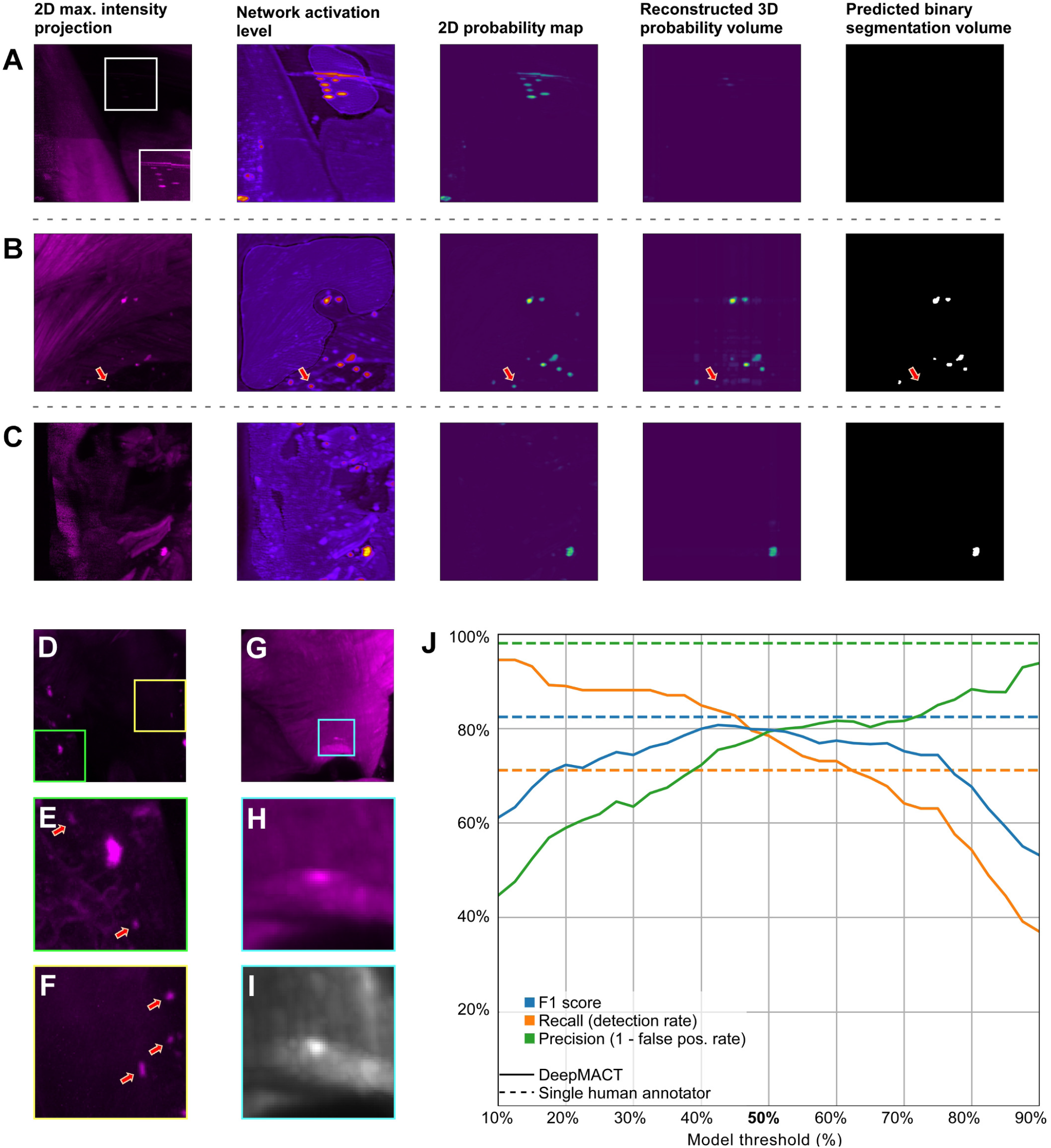
Performance of DeepMACT. **(A-C)** Visualization of the computational stages of DeepMACT for three different regions. **(A)** DeepMACT is capable of identifying very low signal peaks but correctly disregards them at the 3D reconstruction stage; the white inlet shows 10-fold increased brightness. **(B)** While most metastases are correctly identified, few small and dim metastases may be obscured by background structures from some perspectives and consequently be removed at the 3D reconstruction stage (red arrow) **(C)** In many cases, even a single 2D probability map may already be sufficient for a correct prediction. **(D-F)** Example of metastases that were missed by humans but found by DeepMACT (red arrows);brightness of (**G**) and (**F**) increased by 200% compared to (D). (**G,H**) Example of false positive prediction by DeepMACT. The image in (I) shows the same region as in (H) but in the autofluorescence channel (excitation: 488 nm) to confirm that the signal peak in (**H**) is not caused by metastatic tissue. (**J**) DeepMACT performance as a function of model confidence threshold (default: 50%) compared to a single human annotator. While the DeepMACT FI score peaks around 40-50%, the threshold can be adjusted to increase recall or precision.

**Figure S5; related to Figure 5.**
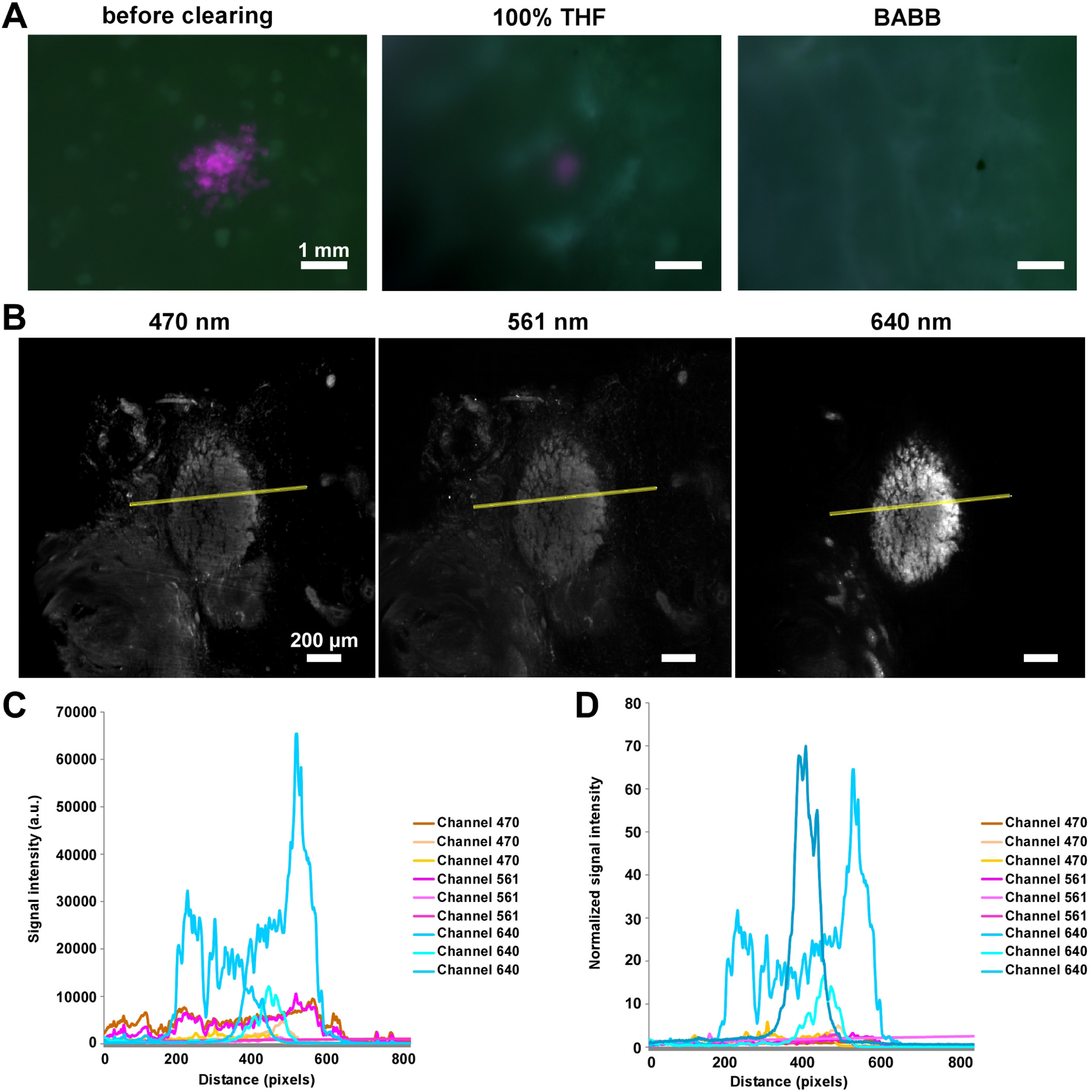
Elimination of endogenously expressed mCherry signal from tumors after vDISCO. **(A)** Tumor metastases in lungs were imaged with a fluorescence stereomicroscope before and after vDISCO clearing, showing that the endogenously expressed mCherry signal was eliminated after THF and BABB incubation steps. **(B)** Light-sheet microscopy images of primary tumor with background imaged in green (ex: 470 nm, left), with mCherry signal in the red (ex: 561 nm, middle) channel, and the boosted signal (Atto647N) in the far-red channel (ex: 640, right). **(C)** Signal intensity profiles along the yellow lines in panel B were plotted: Channel 470 (orange), Channel 561 (magenta) and Channel 640 (cyan) (n=3 mice). (**D**) Normalized fluorescence signal profiles of the data in (**C**), showing that after vDISCO clearing the endogenous mCherry signal was depleted to background levels.

**Figure S6; related to Figure 5.**
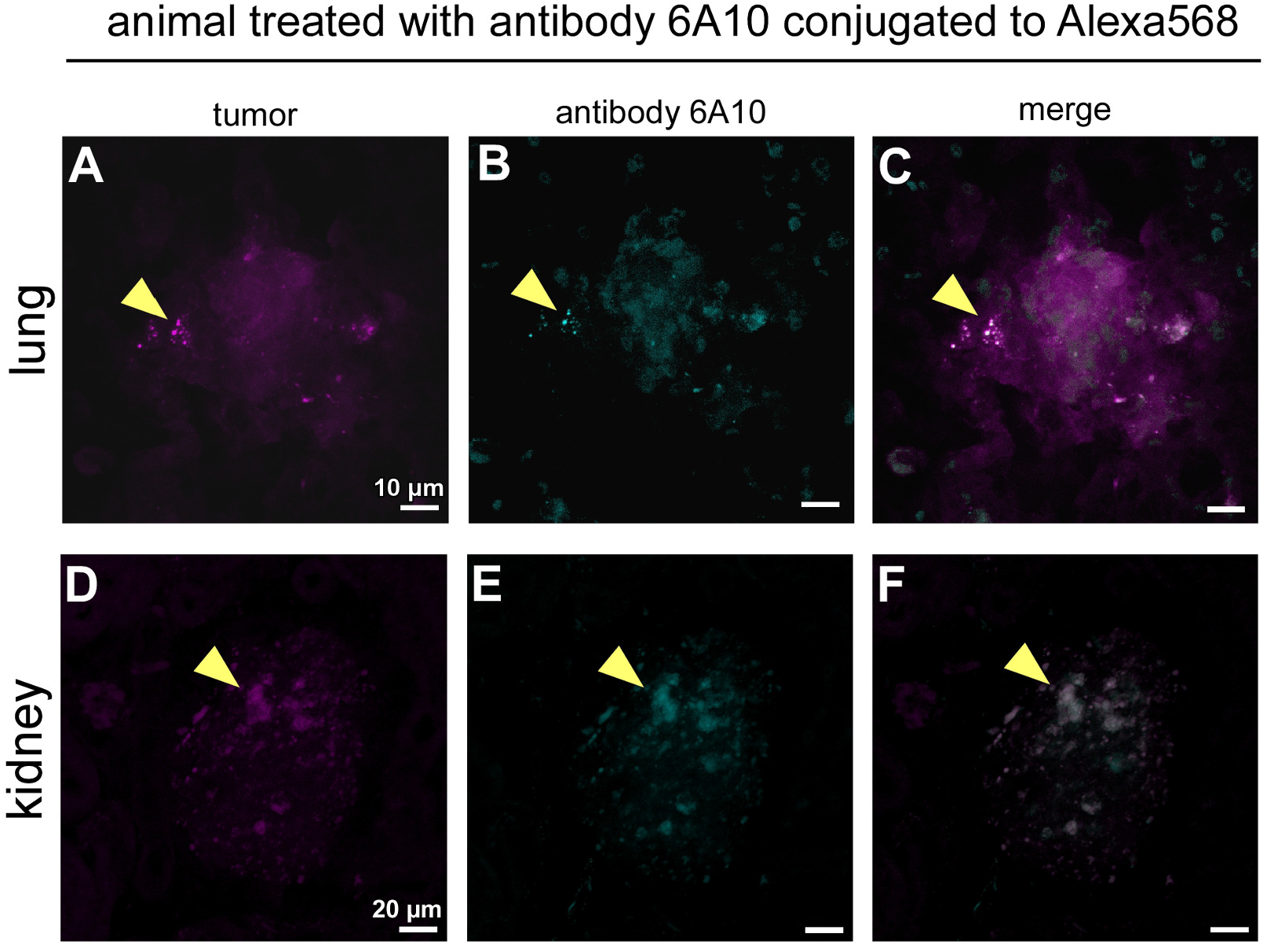
Verification of antibody targeting in different organs by confocal microscopy. **(A-F)** Confocal images of metastases in the lung **(A-C)** and kidney **(D-F)** of an animal labeled with an anti-mCherry nanobody conjugated to Atto647 (magenta) and treated with therapeutic antibody 6A10 conjugated to A1exa568 (cyan). The colocalization of the metastases with the 6A10 antibody is indicated with yellow arrowheads (**C** and **F**).

